# Human DCM-time machine unravels cell state changes during primitive gut tube differentiation

**DOI:** 10.1101/2025.09.26.678757

**Authors:** Marieke E. van Leeuwen, Beatrice F. Tan, Evelyne Wassenaar, Esther Sleddens-Linkels, Ruben Boers, Cristina Gontan, Joost Gribnau

## Abstract

Cell state changes in development and differentiation are directed by gene and enhancer activity changes, the dynamics of which are difficult to study in real time. To be able to define lineage paths and monitor time resolved cell state changes during pluripotent stem cell differentiation, we introduced the DCM-time machine technology (DCM-TM) into human induced pluripotent stem cells (iPSCs). We demonstrate that, at time of induction, human DCM-TM labels active genes and enhancers. In addition, we find that the majority of the DCM methylation labels are propagated during S-phase, which makes it well suited to trace gene and enhancer activity during cell state changes. As proof of concept, we applied the DCM-time machine to study differentiation from iPSCs towards definitive endoderm and primitive gut tube cells. In a comparative analysis with scRNA-seq, we show the capacity of the system to label gene and enhancer activity during differentiation and trace back gene and enhancer activity over time across multiple cell divisions. In addition, we demonstrate that combining DCMTM with methylated DNA sequencing (MeD-seq) enables the detection of CpG methylation changes, which can be correlated with gene and enhancer activity dynamics. Human DCM-TM provides a novel genome-wide lineage tracing tool for iPSCs and will be a significant contribution to the understanding of healthy and pathogenic embryonic development.

## Introduction

Human embryonic development is a remarkable and intricate process that generates more than 300 distinct cell types constituting the human body [1]. A pivotal event in this journey occurs around day 14 of gestation, when gastrulation begins. Gastrulation marks a fundamental phase in embryogenesis, during which the three germ layers, ectoderm, mesoderm, and endoderm, are specified. Prior to this stage, epiblast cells possess the potential to differentiate into any cell type in the body. However, with the onset of gastrulation, these cells commit to a specific germ layer fate, shaping the foundation for future tissue and organ development [2].

Among the germ layers, the endoderm gives rise to vital internal organs, including the entire gastrointestinal and respiratory tracts. Proper development of endodermal tissues is essential for maintaining critical homeostatic functions such as nutrient exchange, digestion, detoxification, and endocrine regulation. Mouse studies indicated that development of the definitive endoderm (DE) is primarily driven by the Activin/Nodal signaling pathway resulting in activation of key transcription factors such as SOX17 and FOXA2 [3, 4]. Further development towards the primitive gut tube requires Wnt and Bone Morphogenetic protein (BMP) signaling, marked by expression of hepatocyte nuclear factor 1 homeobox A (HNF1A) and HNF1B [5, 6, 7].

Dysregulation of these processes can lead to significant physiological dysfunctions [8]. Yet, despite their importance, this crucial period of human embryogenesis remains difficult to study due to both practical and ethical constraints. To overcome these limitations, many in vitro models have been developed to recapitulate endoderm development and development of its derivatives. These include models that recapitulate development of the DE [9] to complex models such as intestinal, stomach, lung and pancreas organoids that mimic organ homeostasis [10, 11, 12, 13]. These models are established using either differentiated induced pluripotent stem cells (iPSCs) or adult stem cells, holding great potential for studying normal and pathological organ development and homeostasis. iPSCs can also be generated from patient-derived somatic cells, enabling the creation of patient-specific models that enhance our ability to investigate disease mechanisms. These in vitro models are essential in understanding the complex molecular processes that direct human embryonic development.

Development involves rapid changes in cell states, driven by shifts in activity of cell signaling pathways that mediate gene regulatory networks. These complex changes at multiple levels of gene regulation are difficult to track in real time. Current studies are usually based on lineage tracing and single-cell RNA sequencing (scRNA-seq) [14, 15]. These techniques depend on in silico assumptions to determine temporal gene expression changes, which can introduce biases. To overcome these limitations, we developed the DCM-time machine (DCM-TM) technology that facilitates whole-genome cell state tracing independent of in silico predictions [16]. DCM-TM utilizes a doxycycline-inducible fusion protein that combines the bacterial methyltransferase DCM with the RNA polymerase II subunit B. Upon induction, this fusion protein labels active gene bodies and enhancers with DCM-specific cytosine methylation at C_me_C(A/T)GG sites. Our studies in mice have demonstrated that DCM methylation does not interfere with transcription and is reliably propagated during DNA replication in S-phase. Therefore, this system enables retrospective tracing of gene and enhancer activity across multiple cell divisions. While mouse models have been invaluable for studying lineage specification, human model systems like iPSCs are better suited for understanding specific human developmental processes. In this study, we introduced DCM-TM into human iPSCs and applied this technology to study iPSC differentiation towards DE and intestinal progenitors. We demonstrate that human DCM-TM (hDCM-TM) efficiently labels gene bodies of active genes and active enhancers, and that the DCM methylation label is efficiently propagated during S-phase. We utilize hDCM-TM to investigate the dynamics in transcription factor networks and gene expression changes during gut tube formation in an in vitro iPSC model system.

## Results

### Human DCM-TM marks active genes in iPSCs with DCM labels

For the development of the human DCM-TM, we built upon the fundamentals established in mice [16]. We coupled the bacterial DCM gene to open reading frame (ORF) encoding human RNA polymerase II subunit B (*POLR2B*) under the control of a doxycycline-inducible (dox) promotor. In addition, we generated a doxycycline-inducible DCM-only construct with a DCM ORF coupled to a nuclear localization signal (NLS) as a control. Using CRISPR/Cas9, these transgenes were introduced into the safe harbor locus *AAVS1* in human iPSCs (Figure S1a). We generated two DCM-*POLR2B* transgenic iPSC lines and one DCM-only transgenic iPSC line and confirmed induction of the DCM-POLR2B fusion protein by PCR and western blot (Figure S1b-c). This analysis also indicated that the DCM-POLR2B fusion protein was expressed at sub-stoichiometric levels compared to wild type POLR2B.

To test whether induction resulted in increased DCM methylation levels, we applied MeD-seq. This technology is based on the DCM and CpG methylation dependent restriction enzyme LpnPI, which generates 32 base pair (bp) reads of fully methylated sites that are sequenced and mapped to the genome (Figure 1a) [17]. DCM-TM iPSCs were harvested after 48 hours of dox induction and isolated DNA was subjected to MeD-seq demonstrating robust induction of DCM methylation (Figure 1b, Table S1). RNA-seq analysis comparing induced versus non-induced iPSCs indicated that expression of only a limited number of genes is affected by the introduction of DCM methylation labels (Figure S1d, Table S2). DCM methylation labels detected with MeD-seq were quantified and normalized based on sequencing depth and number of DCM sites per gene. Correlation analysis comparing these DCM methylation scores with RNA-seq data indicates a very low correlation in uninduced iPSCs (*r* = 0.15, Figure 1c). In contrast, upon activation of the DCM-TM system, a strong correlation was detected (*r* = 0.60, Figure 1c). Moreover, the DCM methylation scores of two independent DCM-TM clones displayed strong inter-clonal correlation post-induction (*r* = 0.98, Figure S1e). DCM methylation was induced at gene bodies of genes known to be active in iPSCs, including *ZFP42, NANOG*, and *SOX2*, and absent in genes not typically active in iPSCs (Figure 1e, S1f). Gene meta-analysis indicated an increase of DCM labels in the gene body accumulating at the transcription start sites (TSS) and transcription end sites (TES), which correlate with gene expression levels in RNA-seq (Figure 1d, S1g). Similar to our findings in mice, the profile plots closely resemble ChIP-seq profiles of RNA polymerase II (Figure S1h). In contrast, iPSCs harboring a DCM-only transgene did not show this pattern after induction (Figure S1i). These results demonstrate that we have generated iPSCs harboring a DCM-*POLR2B* fusion gene that, upon induction, efficiently methylates gene bodies of active genes.

**Figure 1.**
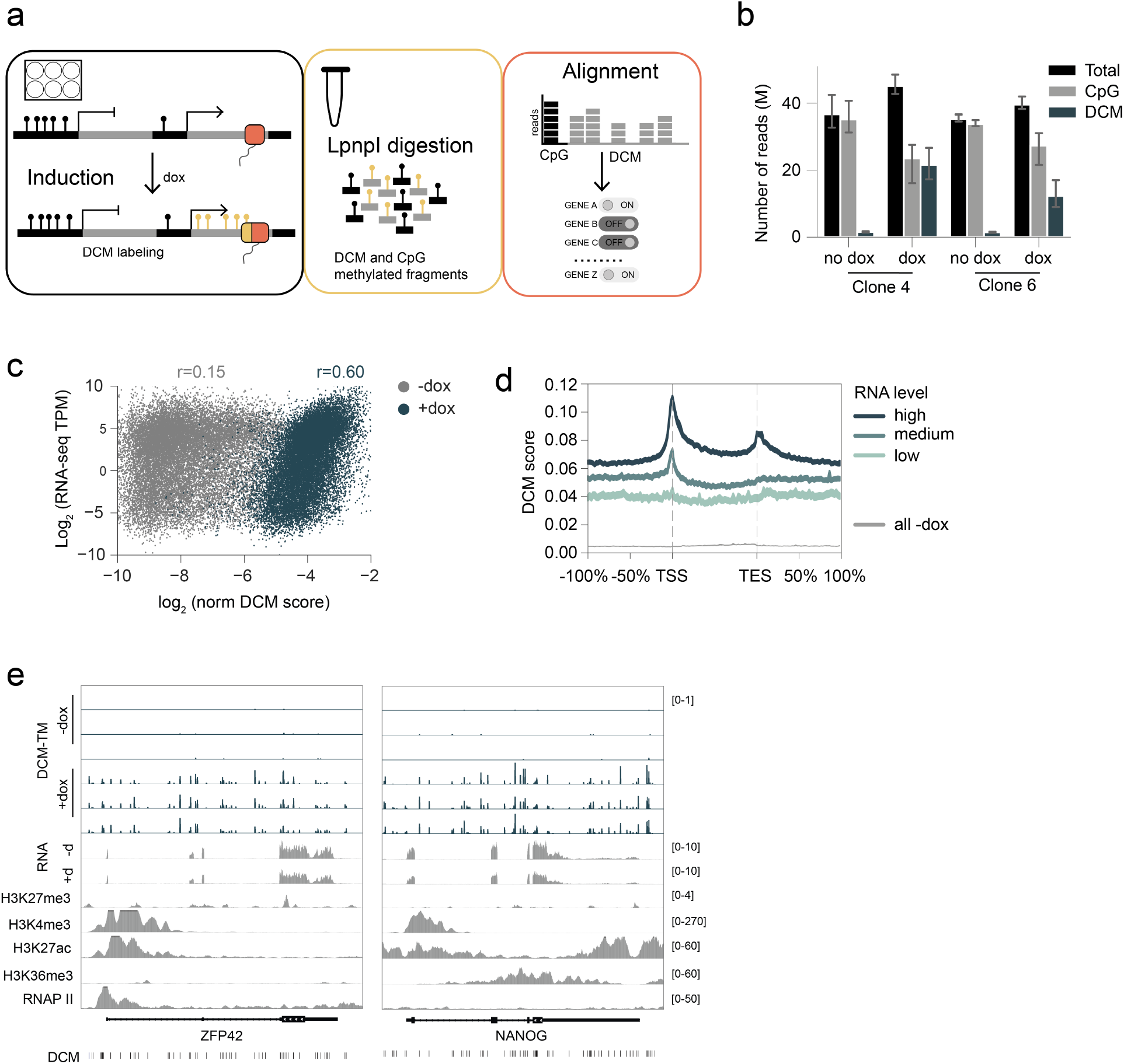
hDCM-TM in iPSCs. **a**. Schematic overview of human DCM-TM and MeD-seq pipelines. Black pins represent CpG methylation and yellow pins represent DCM methylation. **b**. Number of sequenced MeD-seq reads, including total, CpG-specific and DCM-specific reads, pre- and post-induction. Mean of *n* = 3 with 95% confidence interval is shown in millions for two separate clones. **c**. Correlation between DCM methylation labels and bulk RNA-seq expression in uninduced (gray) and dox-induced (blue) samples of DCM-TM clone 4. Pearson correlation coefficients (r) are indicated. **d**. Profile plots of DCM methylation labelling in DCM-*POLR2B* iPSCs +dox, showing distribution of DCM reads along the gene body, upstream of the transcription start site (TSS, −100% of the gene length), and downstream of the transcription end site (TES, +100%). Genes are split in categories based on gene expression levels in bulk RNA-seq data. The gray profile plot shows the DCM methylation levels in −dox iPSCs across all genes. Mean DCM scores with 95% confidence intervals are shown. **e**. Genome browser view showing DCM scores, RNA-seq expression and ChIP-seq enrichment at loci of pluripotency genes *ZFP42* and *NANOG* in dox-induced DCM-*POLR2B* iPSCs. The RNA-seq track shows the mean expression of *n* = 3. d, doxycycline. RNAP II: RNA polymerase II.

### Efficient propagation of the DCM label in S-phase

To enable tracing of cell state changes, it is essential that the DCM labels are propagated during S-phase. To establish the propagation rate of DCM methylation in DCM-TM iPSCs, we performed pulse-chase experiments where we tracked both the cell division rate as well as the DCM methylation loss in the same cell (Figure 2a). To do so, we induced undifferentiated iPSCs for 48 hours with doxycycline and subsequently labeled the cells with CellTrace, which is a permanent fluorescent label that is equally diluted every cell division. We determined the propagation rate at 24, 48 and 72-hour intervals in a pulse-chase experiment by comparing the decrease in the DCM/CpG ratio to the number of cell divisions detected by the dilution of the CellTrace marker over the same interval (Figure 2b, S2a-c, Table S1). These results indicate an average propagation rate of 72% per cell division, suggesting that, based on in silico dilution simulations, active genes can be traced for up to 13 cell divisions (Figure 2c). Importantly, propagation and loss of DCM label appeared consistent in different genomic regions (Figure 2d). Our findings confirm that DCM methylation labels are propagated at an efficiency that facilitates tracing of gene activity over many cell divisions.

**Figure 2.**
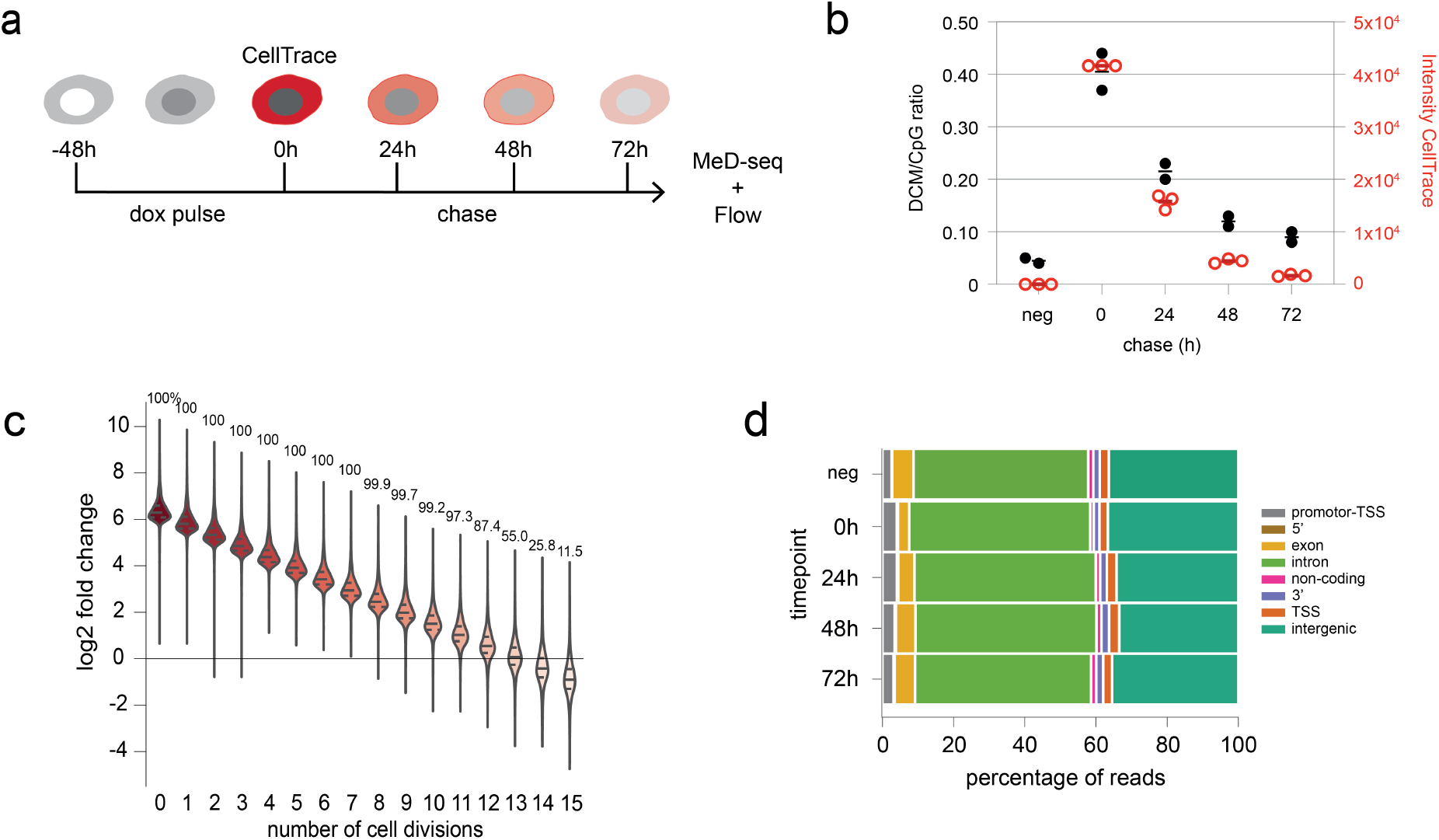
Propagation of DCM methylation in iPSCs. **a**. Experimental overview of the DCM propagation experiment in DCM-TM iPSCs. **b**. DCM/CpG ratio (*n* = 2) and median CellTrace intensity (*n* = 3) during the pulse-chase experiment in **a. c**.In silico prediction of active gene detection across successive cell divisions in a chase experiment. Log_2_ fold change between −dox samples and in silico diluted +dox samples are shown. Above each violin, the percentage of active genes with expected DCM levels exceeding those of the −dox samples is indicated. **d**. Genomic annotation of DCM methylation reads from the pulse-chase experiment shown in **a**.

### Assessment of primitive gut tube differentiation using the hDCM-TM

We applied the hDCM-TM to investigate cell state changes in iPSC differentiation towards DE and subsequently into primitive gut tube (PGT) cells using the first two phases of the STEMdiff pancreatic progenitor kit.

Differentiation of iPSCs was highly efficient, as validated by RT-qPCR for iPSC, DE and PGT marker genes and flow analysis for DE marker proteins CXCR4 and c-KIT (Figure S3a-b). To test whether the DCM-*POLR2B* transgene is efficiently induced in all progenitor cell types, we induced DCM-TM iPSCs with doxycycline during the last 24 hours of each differentiation stage. In addition, DCM-TM iPSCs were induced for 24 hours followed by a 72-hour chase period and harvested at the next differentiation stage (chase) (Figure 3a). MeD-seq analysis of isolated DNA indicated that all samples showed increased DCM methylation labelling levels compared to non-induced iPSCs (Figure S3c, Table S1). Comparison of pulse-only and pulse-chase samples indicated loss in DCM methylation that can be attributed to propagation of the DCM label. To account for this, counts were normalized to DCM scores by normalizing for the number of DCM sites and sequencing depth, followed by scaling to the same sum of genic normalized values across all samples (see Methods). Profile plots of DE and PGT samples displayed patterns similar to those of iPSCs showing enrichment of DCM methylation at the TSS and within the gene body (Figure S3d-e). This demonstrates that we successfully generated a high-quality MeD-seq dataset, capturing DCM methylation profiles for each distinct cell type in both their present and past states.

**Figure 3.**
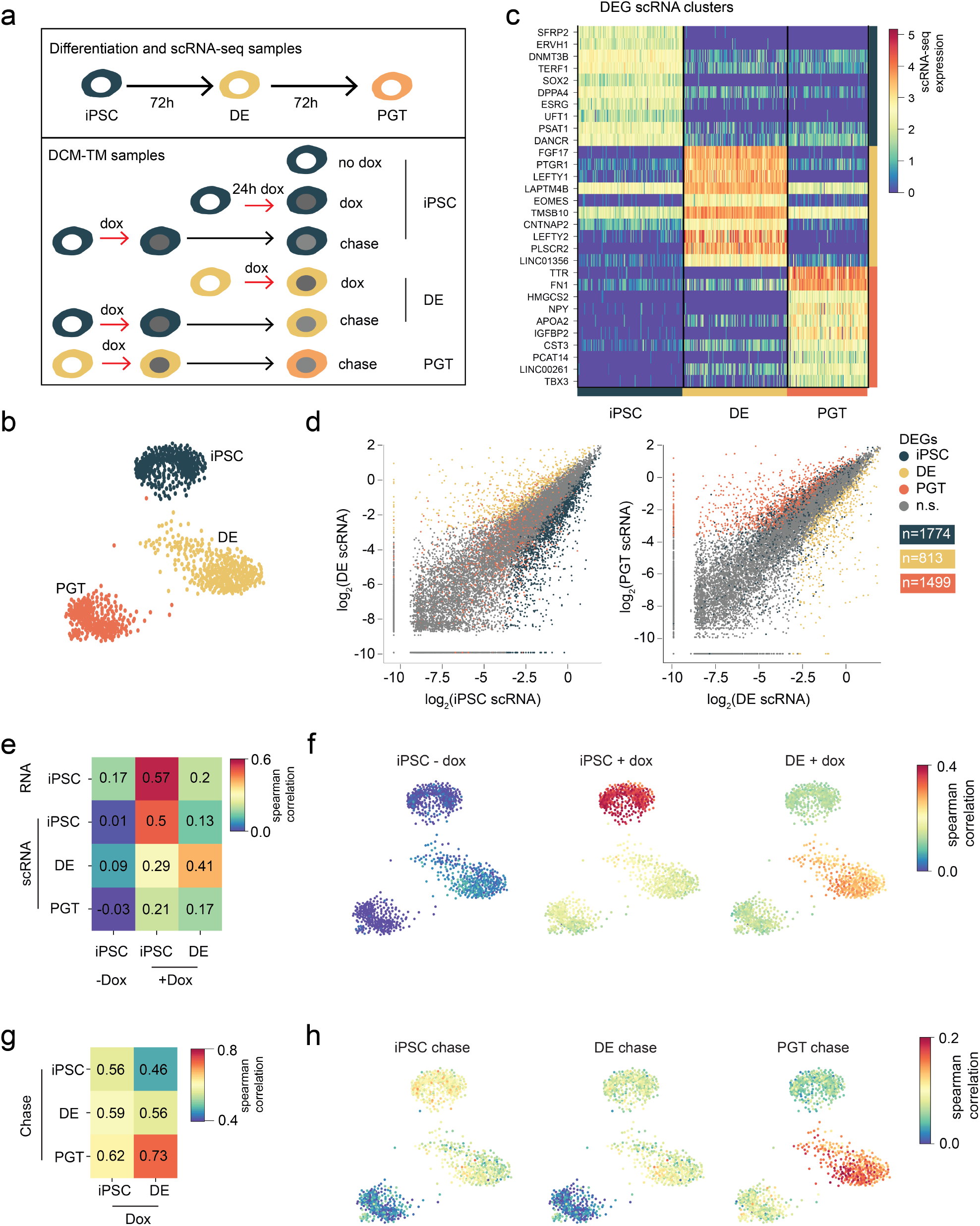
Genome-wide analysis of primitive gut tube differentiation. **a**. Schematic overview of experimental procedures. scRNA-seq was performed for each cell state, and MeD-seq was conducted both immediately after 24-hour dox induction (red arrow) and following a chase period of 72 hours (black arrow). **b**. Force-directed layout of scRNA-seq data generated by Harmony, colored by cell type. **c**. Heatmap showing expression of top 10 DEGs from each cell type in the scRNA-seq dataset. **d**. Scatter plots showing correlation between pseudobulk expression data of scRNA-seq cell types (log2 of normalized counts), with DEGs per cell type highlighted (iPSCs *n* = 1,774, DE *n* = 813 and PGT *n* = 1,499). **e**. Heatmap showing Spearman correlation between RNA-seq, scRNA-seq pseudobulk and DCM +dox and −dox samples, calculated using only the scRNA-seq DEGs shown in **d. f**. Spearman correlation between normalized scRNA-seq counts per cell and DCM methylation scores in iPSC^−dox^ (left), iPSC^dox^ (center), and DE^dox^ (right) samples, mapped onto scRNA-seq layout in **b**. Correlation coefficients were calculated using the scRNA-seq DEGs shown in **d. g**. Heatmap showing Spearman correlation between DCM dox pulse and chase samples, calculated using only the scRNA-seq DEGs shown in **d. h**. Spearman correlation between normalized scRNA-seq counts per cell and DCM methylation scores in iPSC^chase^ (left), DE^chase^ (center) and PGT^chase^ (right) samples, mapped onto scRNA-seq layout in **b**. Correlation coefficients were calculated using the scRNA-seq DEGs shown in **d**.

As a reference dataset, we generated a scRNA-seq gene expression map of cells in the iPSC, DE and PGT states using SORT-seq. Individual cells from the three differentiation stages formed distinct clusters, as visualized by a force-directed layout generated using Harmony [8] (Figure 3b). Differential gene expression analysis between these clusters revealed many marker genes among the differential expressed genes (DEGs), including *SOX2, EOMES* and *APOA2* for iPSC, DE and PGT, respectively (Figure 3c, Table S3). Next, we generated pseudobulk gene expression profiles for each cluster. Even though DEGs between cell types were detected, the overall gene expression between cell types displayed a high correlation (*r* = 0.91, Figure 3d, S4a). This high similarity reflects the fact that the studied cell types are all at a very early stage of development and are closely related in the differentiation process. Therefore, we validated the DCM-TM dataset focusing on cell type-specific differences using correlation analyses on the DEGs detected by scRNA-seq. This analysis robustly demonstrates that DCM methylation scores of the dox samples correlate best with scRNA-seq expression of their respective cell type (Figure 3e-f, S4b). For instance, iPSC^dox^ has the strongest correlation with both bulk RNA and scRNA-seq data of iPSCs, while DE^dox^ correlates best with DE cluster in the scRNA-seq data. Neither of these cell states showed a strong correlation with the scRNA-seq data of PGT cells (Figure 3e-f, S4c). In contrast, when analyzing the chase samples, the DCM methylation labels reflected the cell state at time of induction, rather than at the time of isolation. For instance, DE^chase^ correlated best with iPSC^dox^, and PGT^chase^ correlated best with DE^dox^ (Figure 3g-h, S4d), which is consistent with their stepwise state transitions. These results demonstrate that the hDCM-TM technology can be efficiently utilized to determine the genome-wide gene expression profiles of a prior cell state in human cells.

### Gene dynamics in primitive gut tube differentiation

To study the gene dynamics underlying cell state transitions, we next focused on DCM methylation scores at the level of individual genes. We began by examining key marker genes detected by scRNA-seq at multiple timepoints of differentiation. As expected, DCM labeling was highest at the corresponding iPSC^dox^ and DE^dox^ samples for iPSC and DE marker genes, respectively. In the pulse-chase samples, DCM labeling of iPSC-specific genes mostly resembled labeling in iPSC^chase^ and DE^chase^, whereas labeling of DE-specific genes more closely resembled that in PGT^chase^ samples (Figure 4a). For instance, pluripotency-associated genes such as *ZFP42, NANOG* and *POU5F1* showed the highest DCM labeling in iPSC^dox^ samples, as well as in the iPSC^chase^ and DE^chase^ samples, with gene activity further supported by scRNA-seq and histone modification data (Figure 4b, S5a-b). Likewise, genes expressed in DE, including *EOMES, GATA4* and *FGF17*, had the highest DCM score in the DE^dox^ and PGT^chase^ samples (Figure 4c, S5c-d). In contrast, for PGT marker gene *TBX3* no such clear pattern was apparent (Figure S5e). We next investigated gene dynamics solely based on DCM methylation profiles, to also uncover dynamic genes not captured by scRNA-seq. Differential methylation analysis between iPSC^dox^ and DE^dox^ cells revealed many differentially DCM methylated genes (DDMs), including 1,714 iPSC DDMs and 1,899 DE DDMs (Figure 4d, Table S4). Many marker genes for iPSC and DE were among the top hits, including *FOXN3* and *GATA4*. The DDMs specific to iPSC and DE showed significant enrichment in the corresponding iPSC and DE clusters of the scRNA-seq data (Figure 4e, S5f). Subsequently, we evaluated the potential of cell state tracing with DCM-TM. We compared the DE^chase^ and PGT^chase^ samples to identify iPSC- and DE-specific genes, respectively, based on DCM labeling. In total, 388 DDMs were upregulated in DE^chase^ and 1,106 DDMs in PGT^chase^ (Figure 4f, Table S5). The number of DDMs was lower than in the dox samples, as expected based on partial loss of DCM labels during propagation. However, when examining the top DEGs, many genes labeled in the dox samples were also detected in the chase samples, including *GATA4, ROR2* and *EGFLAM* as DE-specific genes. Enrichment of the DDMs in the chase samples with the scRNA-seq dataset confirmed that iPSC- and DE-specific genes were indeed labeled by DCM-TM (Figure 4g, S5g). These findings demonstrate that human DCM-TM can capture gene dynamics both at induction and retrospectively over time, making it a powerful system suitable for cell state tracing experiments.

**Figure 4.**
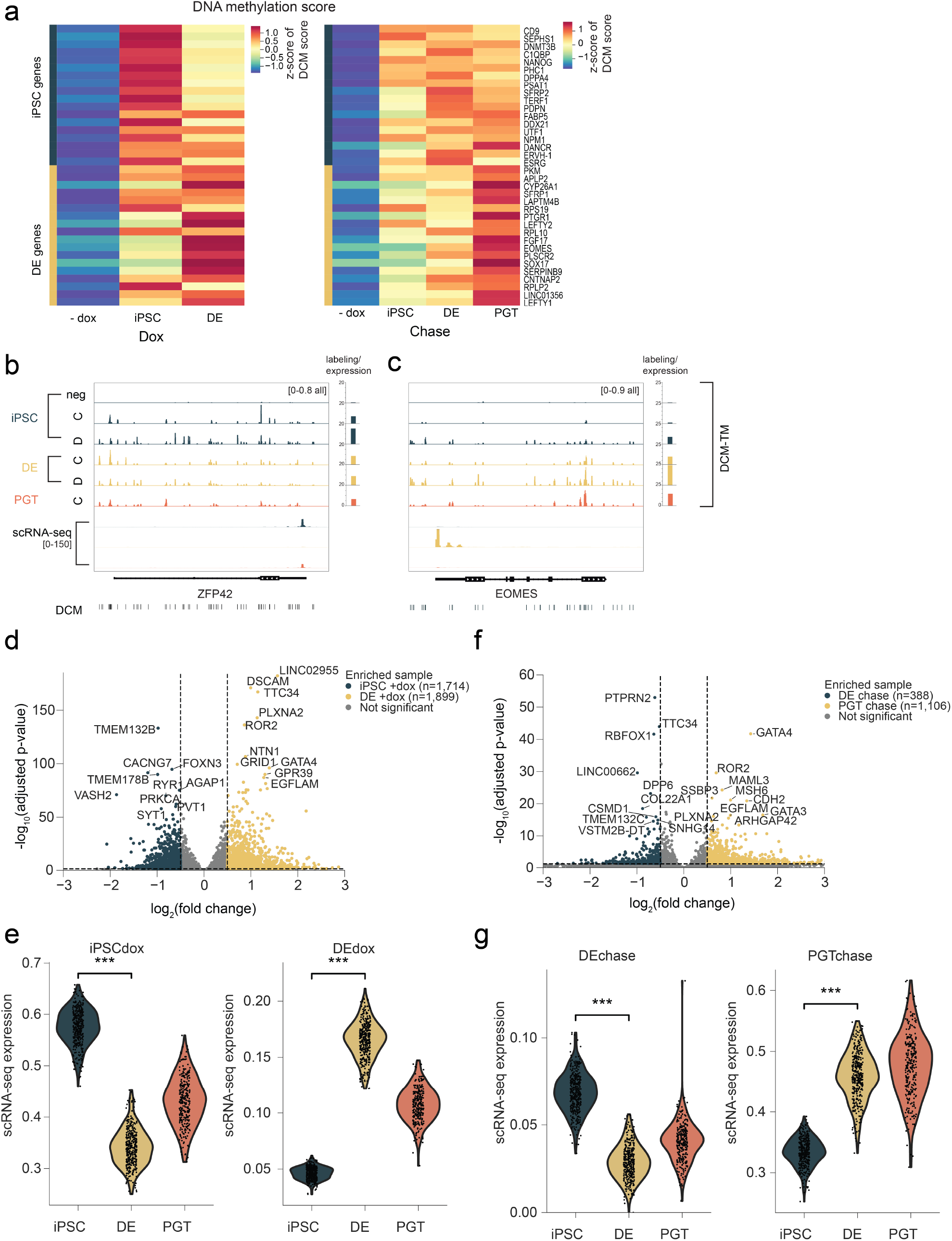
Gene dynamics in primitive gut tube differentiation. **a**. DCM labelling of top DEGs in the scRNA-seq dataset. Z-scores of average DCM scores (*n* = 3) per condition are shown, calculated separately for the dox (left) and chase (right) samples. **b**,**c**. Genome browser view at loci of *ZFP42* (pluripotency marker, **b**) and *EOMES* (DE marker, **c**). Tracks show DCM methylation (average of *n* = 3) and scRNA-seq pseudobulk tracks of iPSC, DE and PGT cells. The barplots on the right quantify all DCM scores overlapping *ZFP42* (**b**) and *EOMES* (**c**) per sample. C, chase; D, dox. **d**,**f**. Differentially DCM-methylated genes (DDMs) between iPSC^dox^ and DE^dox^ (**d**) and between DE^chase^ and PGT^chase^ (**f**). Significant DDMs are highlighted in blue and yellow. **e**,**g**. Average expression of DDM genes identified in **d** and **f** across different cell types from the scRNA-seq dataset. *P* values were calculated using Mann-Whitney tests (*** *P* value < 0.001).

### Enhancer activity during differentiation

Enhancers play a central role in gene regulatory networks by boosting the transcription of specific genes, thereby ensuring precise spatial and temporal gene expression essential for proper differentiation. Analysis of H3K27ac ChIP-seq data in iPSCs [18] and DE [19] revealed that many H3K27ac peaks showed enrichment of DCM labeling in the dox-treated samples, in contrast to the uninduced samples (Figure S6a-b). This indicated that the DCM-TM system can be used to detect active enhancers, as shown previously for DCM-TM in mice [16]. DCM labeled enhancers were called using a sliding window approach, interrogating all intergenic DCM sites >1kb from TSS, followed by a differential test between two cell states and merging of overlapping significant regions. Differential DCM methylation analysis between iPSC^dox^ and DE^dox^ identified enhancer sites at intergenic regions and proximate to known marker genes *SOX2* and *GATA4* (Figure 5a-b, S6c-d) [20]. In total, we identified 3,949 iPSC-specific and 9,623 DE-specific enhancers (Figure 5c, S6e-f). Since this analysis focuses on changes in enhancer activity between iPSC and DE, rather than all active enhancers marked by H3K27ac, not all H3K27ac peaks overlap with DCM-labelled enhancers (Figure S6c-d). Active iPSC and DE-linked enhancers were also labelled in the corresponding chase samples (Figure S6e-f), indicating that enhancer activity can be traced over time.

**Figure 5.**
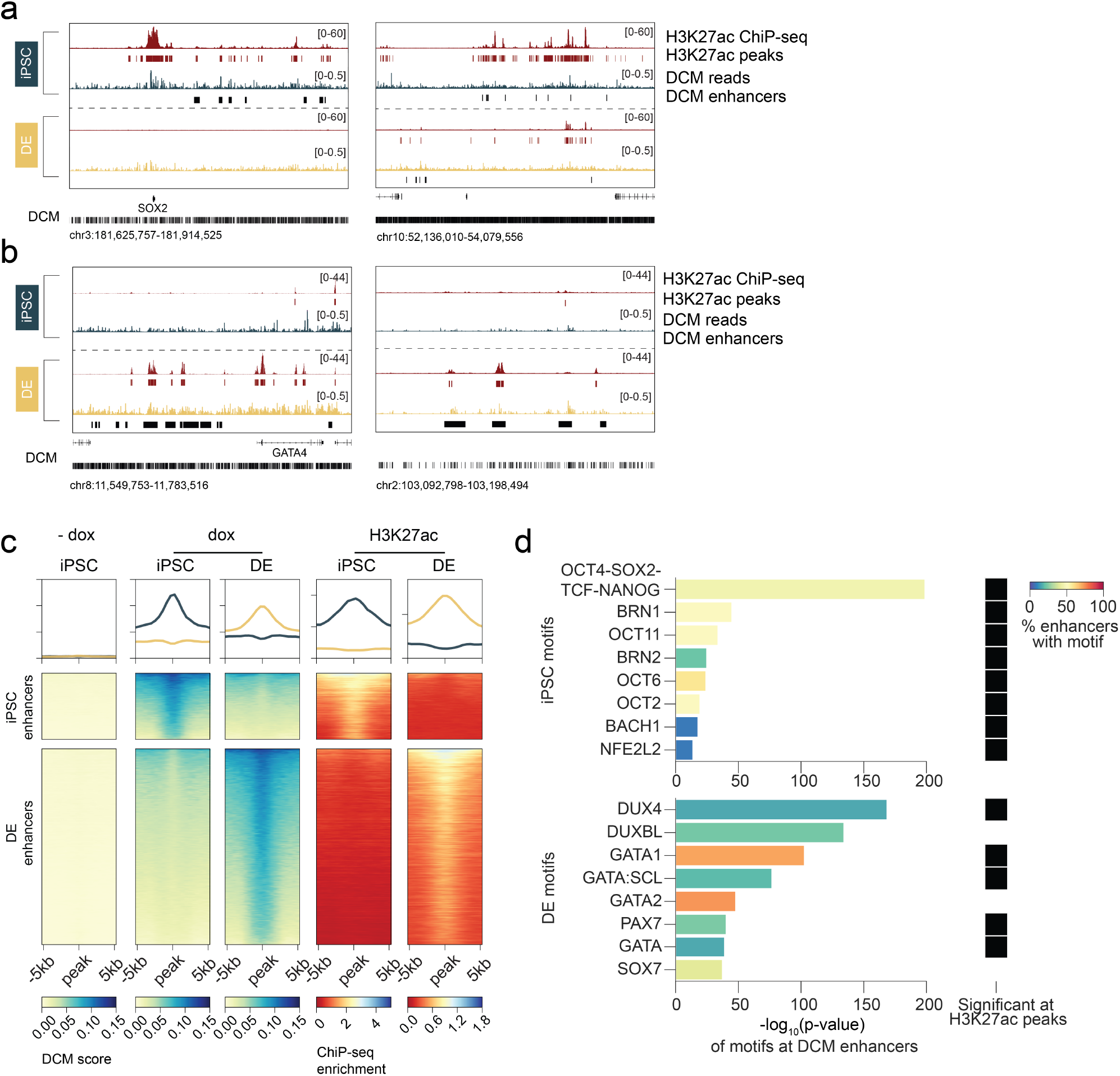
Enhancer regions are labeled with DCM methylation. **a**. Genome browser view of DCM-labelled enhancer regions, including iPSC enhancers upstream of *SOX2* (left) and both iPSC- and DE-specific enhancers on chromosome 10 (right). Tracks show iPSC and DE H3K27ac ChIP-seq enrichment and corresponding significant peaks (in red), and DCM scores of iPSC^dox^ and DE^dox^ (average of *n* = 3) with significant DCM-labelled enhancers below (in blue and yellow, respectively). Note that only intergenic enhancers are called from the DCM data, meaning no enhancers overlapping with SOX2 are identified. **b**. Genome browser view of DCM-labelled enhancer regions, including DE enhancers upstream of *GATA4* (left) and in an intergenic region on chromosome 2 (right). Tracks shown correspond to those in **a. c**. Heatmaps showing enrichment of DCM scores in iPSC^−dox^, iPSC^dox^ and DE^dox^, as well as H3K27ac ChIP-seq datasets from iPSC and DE, at DCM-labelled enhancers specific to iPSCs (top) and DE (bottom). Profile plots have the same y-axis range as the corresponding heatmap. **d**. Significantly enriched motifs in iPSC (top) and DE (bottom) enhancer regions. Barplots show significance values reported by HOMER, with bars colored by the percentage of enhancers containing the motif sequence. Black boxes indicate whether the motif is significantly enriched in H3K27ac peaks of the corresponding cell type.

Motif analyses on the enhancer regions was performed to predict transcription factor (TF) binding during DE differentiation (Figure 5d). In addition, analysis of motifs enriched in H3K27ac peaks in iPSCs and DE revealed that many motifs were significantly enriched in both datasets. In iPSC enhancer regions, the OCT4-SOX2-TCF-NANOG motif was enriched, consistent with previous findings [21]. Moreover, several OCT TF motifs, including BRN1/2 and OCT2/6/11, were enriched in these regions, aligning with their known roles in maintaining iPSC identity and pluripotency [22, 23]. In contrast, DE enhancers showed an enrichment of TF binding motifs from the GATA-family, which have been reported as regulators of endoderm formation [24, 25, 26]. This demonstrated the capacity of the hDCM-TM to detect active enhancers simultaneously with active genes, which can be used to predict key TF activity during differentiation.

### Large changes in CpG methylation occur early in lineage determination

Because MeD-seq, used to detect DCM methylation labels, also captures CpG methylation genome-wide, we also examined changes in CpG methylation during differentiation. Since CpG methylation is unaffected by the DCM-TM, all previously obtained MeD-seq samples could be used for this analysis, regardless of the timing of dox induction. We analyzed changes in CpG methylation in two steps, from iPSC to DE and subsequently from DE to PGT. Differential methylation analysis identified 9,730 differentially methylated regions (DMRs) between iPSCs and DE (Figure 6a) and 1,757 DMRs between DE and PGT cells (Figure 6b). These findings suggest that the largest differences in the epigenome occur early in differentiation. Categorizations of the DMRs showed that most extensive changes occur at CpG islands, predominantly with an increase in methylation (Figure 6c-d). The emphasis on hypermethylation indicates that as cells commit to a differentiation pathway, their epigenome gets increasingly restrictive. CpG methylation at promoter regions further supported this, revealing a shift towards gene silencing, generally characterized by hypermethylation of the promoter.

**Figure 6.**
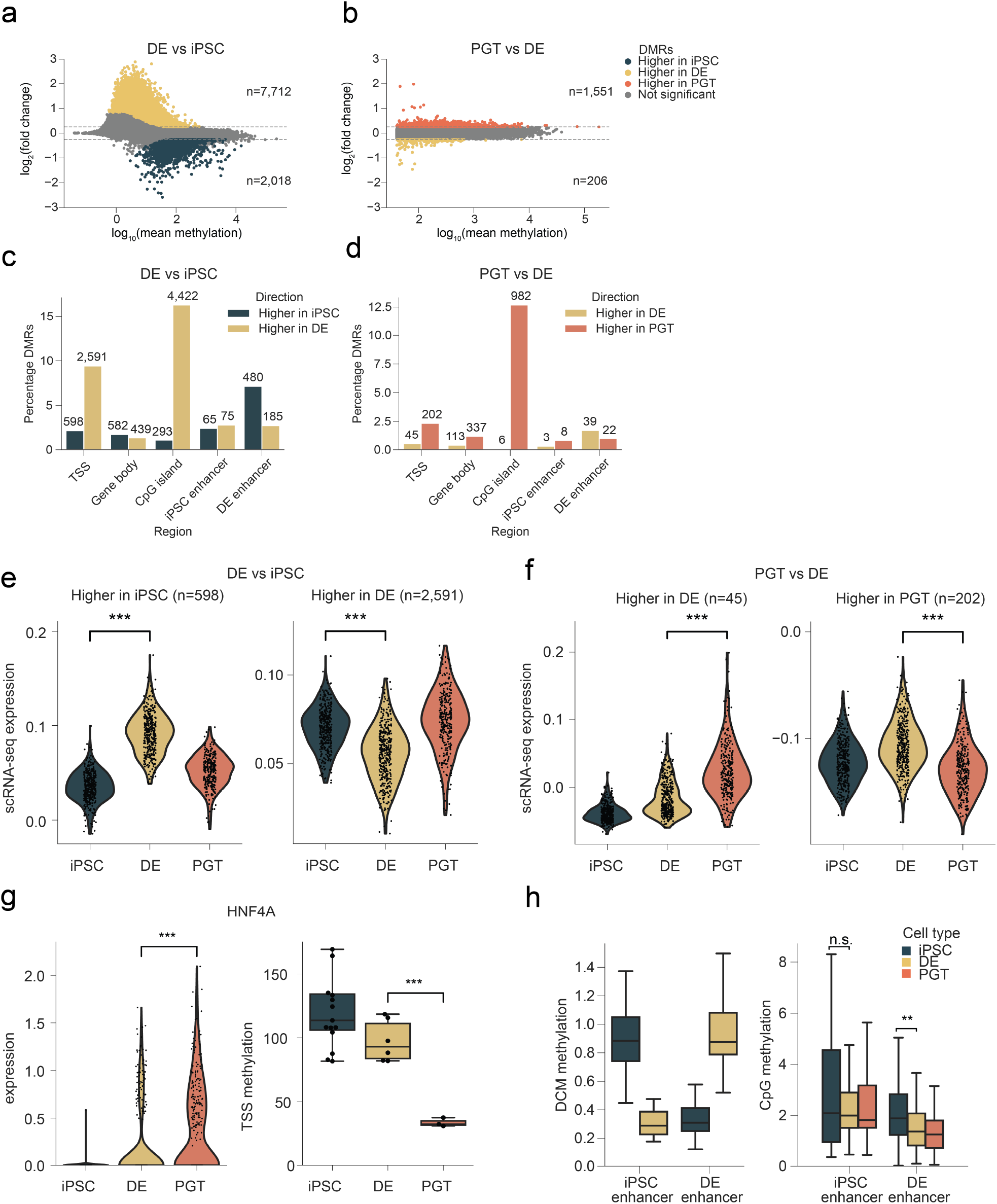
Enhancer regions are labeled with DCM methylation. **a**,**b**. MA plot of differentially CpG-methylated regions (DMRs) between DE and iPSC cells (**a**) and between PGT and DE cells (**b**). Significant DMRs are highlighted in blue, yellow and orange for hypermethylation in iPSCs, DE and PGT cells, respectively. **c**,**d**. Barplot showing percentages of DMRs found in (**a**,**b**), divided by genomic region. For each category, the percentage of significant regions are shown with the absolute numbers of DMRs above the bar. TSS, transcription start site. **e**,**f**. Violin plots showing scRNA-seq gene expression of genes with differentially methylated TSSs, split by cell type. **g**. scRNA-seq expression of HNF4α (left) and CpG methylation levels at the TSS (right), split by cell type. The box plots display the median (black line), the interquartile range (box limits) and 1.5x of the interquartile range (whiskers). **h**. Cell type-specific DCM scores and CpG methylation levels at DCM enhancers with significant CpG methylation changes between iPSC and DE samples. Enhancers were split by cell type with the most DCM labelling. The box plots display the median (black line), the interquartile range (box limits) and 1.5x of the interquartile range (whiskers). *P* values were calculated using Mann-Whitney tests (* *P* < 0.05, ** *P* < 0.01, *** *P* < 0.001).

In the transition from iPSCs to DE, 598 promoter regions lost CpG methylation, while 2,591 promoters became hypermethylated in DE (Figure 6c, S7a). Examination of genes with significant promoter DMRs showed that these CpG changes correlate negatively with expression changes, as previously described [27, 28] (Figure 6e, S7b). For example, *OCT6*, an active TF in iPSCs, displayed both significant promoter hypomethylation and significantly higher gene expression levels in iPSCs compared to DE (Figure S7c, S6c). In the cell state change from DE to PGT, 45 promoters were hypomethylated, while 202 promoters were hypermethylated in PGT cells (Figure S7d). These changes were significantly linked to up- and downregulation of gene expression, respectively (Figure 6f, S7e). Notably, the hepatocyte nuclear factor *HNF4A*, a key transcription factor for PGT development, is upregulated in PGT cells, while its promoter becomes hypomethylated (Figure 6g).

To assess the role of CpG methylation in enhancer regulation, we identified significant hypo- and hyper-methylation events at the enhancer regions identified from the DCM labelling dataset (Figure 5c). In total, 805 enhancers exhibited significant CpG methylation changes during the transition from iPSCs to DE (Figure 6c), whereas only 72 enhancers showed changes between DE and PGT (Figure 6d). It is important to note that the DCM-TM dataset did not include samples that would allow identification of PGT-specific enhancers, and therefore, these regions were not analyzed. Overall, enhancer regions were more frequently hypomethylated than hypermethylated during differentiation, in contrast to the methylation dynamics observed at promoter regions and CpG islands (Figure 6c-d). While CpG methylation dynamics at iPSC enhancers is very limited between cell states, DE enhancers displayed a significant shift towards hypomethylation compared to iPSCs (Figure 6h). This pattern suggests a positive correlation between enhancer activity and loss of CpG methylation at enhancers regulating DE differentiation. Together, these findings demonstrate that DCM-TM in combination with MeD-seq provides a powerful system for studying both active genes and enhancers across different cellular states, while simultaneously capturing the corresponding CpG methylation landscape. The observed correlations between gene expression, enhancer activity and CpG methylation highlight that cell state transitions are regulated through multiple layers of epigenetic and transcription control.

## Discussion

In this study we have developed the hDCM-TM, a novel whole-genome cell state tracing technique which enables retrospective analysis of gene and enhancer activity in human pluripotent cells. We applied this technology to investigate differentiation of iPSC to DE and PGT, uncovering dynamic changes in gene expression during differentiation. In addition, we revealed enhancer activity dynamics and related this to transcription factor network changes. We, furthermore, demonstrate the simultaneous detection of DCM and CpG methylation, facilitating combined analysis of gene and enhancer activity changes in time in relation to the establishment of the epigenome.

Our study shows that the human DCM-POLR2B fusion is expressed at low levels compared to wild type POLR2B, and the high ratio between gene body labeling of active and inactive genes after induction of hDCM-TM indicates that most of the fusion protein generated is integrated in the RNA polymerase 2 complex. As we reported previously in mice, comparison of DCM label accumulation with RNA-seq and histone modifications shows that DCM-TM efficiently labels gene bodies of active genes as well as active enhancers genome-wide. The fact that highly expressed genes accumulate more DCM labels compared to lowly expressed genes is probably related to the inefficient labeling by the machinery, as we showed before for the murine DCM-TM system [16].

We demonstrate that, in human cells, gene and enhancer activity can be traced over multiple cell divisions and utilized to investigate TF dynamics in relation to cell state changes. An obstacle in the application of the hDCM-TM is the propagation efficiency of the DCM methylation label. Currently, there is a limit to the number of cell divisions that can be investigated and an in silico correction for the loss of DCM methylation was performed. In addition, our data showed a modest overlap between the chase samples and the state used as the viewpoint. This could be explained by prolonged activity of the DCM-POLR2B fusion protein labeling genes during the transition to the next state.

Our study revealed distinct differences in genome-wide CpG methylation changes during differentiation. In the initial transition from iPSCs to DE cells, we observed substantial and highly robust alterations in CpG methylation patterns. In contrast, the subsequent differentiation step from DE to PGT cells exhibited comparatively smaller changes. This pattern aligns with in vivo findings in mice, where CpG methylation dynamics peak immediately after implantation and during early gastrulation [29, 30]. Notably, these changes primarily involved increased CpG methylation associated with gene repression, most prominently at gene regulatory elements during the transitions from iPSC to DE and from DE to PGT. This finding suggests that as iPSCs differentiate, the epigenome becomes progressively more restrictive, likely to ensure the stability of lineage fate decisions in descendant cells.

In summary, we generated a system that enables labeling of active genes and enhancers that can be traced back in time and related to epigenomic changes. Therefore, hDCM-TM provides a unique platform to investigate and optimize iPSC differentiation protocols by isolating properly differentiated cells and retrospectively analyzing cell state changes to determine the correct signals at the appropriate time during differentiation. hDCM-TM also provides new opportunities to study the effect of pathogenic mutations and will aid in retrospective determination of cell states that are sensitive or insensitive to treatments.

## Methods

### Cell lines

All cells were maintained in 5% CO2 incubators at 37 °C and were routinely tested for mycoplasma contamination and karyotyped. Before and after nucleofection, human iPSC lines were genotyped using the Infinium Global Screening Array (GSA) (Illumina) to ensure genome stability. All experiments were performed using the WTC-11 human male iPSC line (Coriell Repository, GM25256). The WTC cell line was maintained in Stemflex medium (Gibco, A3349401) on Geltrex™ LDEV-Free Reduced Growth Factor Basement Membrane Matrix (Gibco, A1413201) coated plates. Cells were passaged at 80-90% confluence using ReLeSR (STEMCELL Technologies, #100-0484). Briefly, culture medium was removed from the dish, 1 mL ReLeSR was added to the well and aspirated after 10 seconds. After a 3 min incubation at 37 °C, 5% CO2 cells were resuspended in 3 mL culture medium and plated at the appropriate density. Cells were frozen in PSC Cryopreservation medium (Gibco, A2644401).

Karyotyping was performed as follows: cells were incubated with KaryoMAX Colcemid (12 µL/mL medium, Gibco, #15210-040) for 20 minutes and were dis-sociated using StemPro Accutase Cell Dissociation Reagent (Gibco, a1110501). Cells were pelleted and resuspended in 5 mL, 75 mM KCl at 37 °C and centrifuged for 5 min at 1100 rpm. Cells were resuspended in 6 mL, 75 mM KCl with 20% fixative (3:1, methanol:acetic acid) and pelleted for 10 min at 1100 rpm. Following primary fixation cells were centrifuged 3 times in 5 mL fixative for 10 min at 1100 rpm. After the final centrifugation step the pellet was kept in 0.5 mL fixative. The suspension was mounted using VECTASHIELD® Antifade Mounting Medium with DAPI (Vector Laboratories, H1200-10) and chromosome number was established with fluorescent microscopy.

### Generation of DCM-POLR2B and DCM-NLS fusion genes

To create the human DCM-time machine cell lines, the coding sequence for the DCM-*POLR2B* fusion gene was placed under the control of an m2rtTA promoter and inserted into the *AAVS1* locus using CRISPR/Cas9. For cell lines containing DCM with a nuclear localization label (NLS), DCM was fused to a SV40 NLS sequence in the same repair plasmid. The DCM fusion genes were generated by a combination of restriction-based cloning and a Gibson Assembly® Cloning Kit (New England Biolabs, E5510S). *POLR2B* coding sequence was amplified by PCR from cDNA of HEK293T cells and DCM was amplified from the plasmid encoding the mouse DCM-time machine [16]. At the C-terminus of the DCM-*POLR2B*, there is a P2A followed by the fluorescent protein mCherry. All PCRs were performed using Phusion® High-Fidelity DNA Polymerase (New England Biolabs, M0530L) following the manufacturer’s protocol. Plasmids were amplified in NEB® 5-alpha Competent E. coli (High Efficiency) (New England Biolabs, C2987I) or T7 Express lysY/Iq Competent E. coli (High Efficiency) (New England Biolabs, C3013I). PCR primers can be found in Table 1. The fusion genes were integrated into a modified version of the plasmid pZP (cHS4)X4 TetON-3XFLAG-tdT CAGG-m2rtTA v2 (Addgene #112669), in which the TdTomato cassette was replaced with a Cherry cassette using a gBlocks gene fragment from Integrated DNA Technologies (IDT).

**Table 1.**
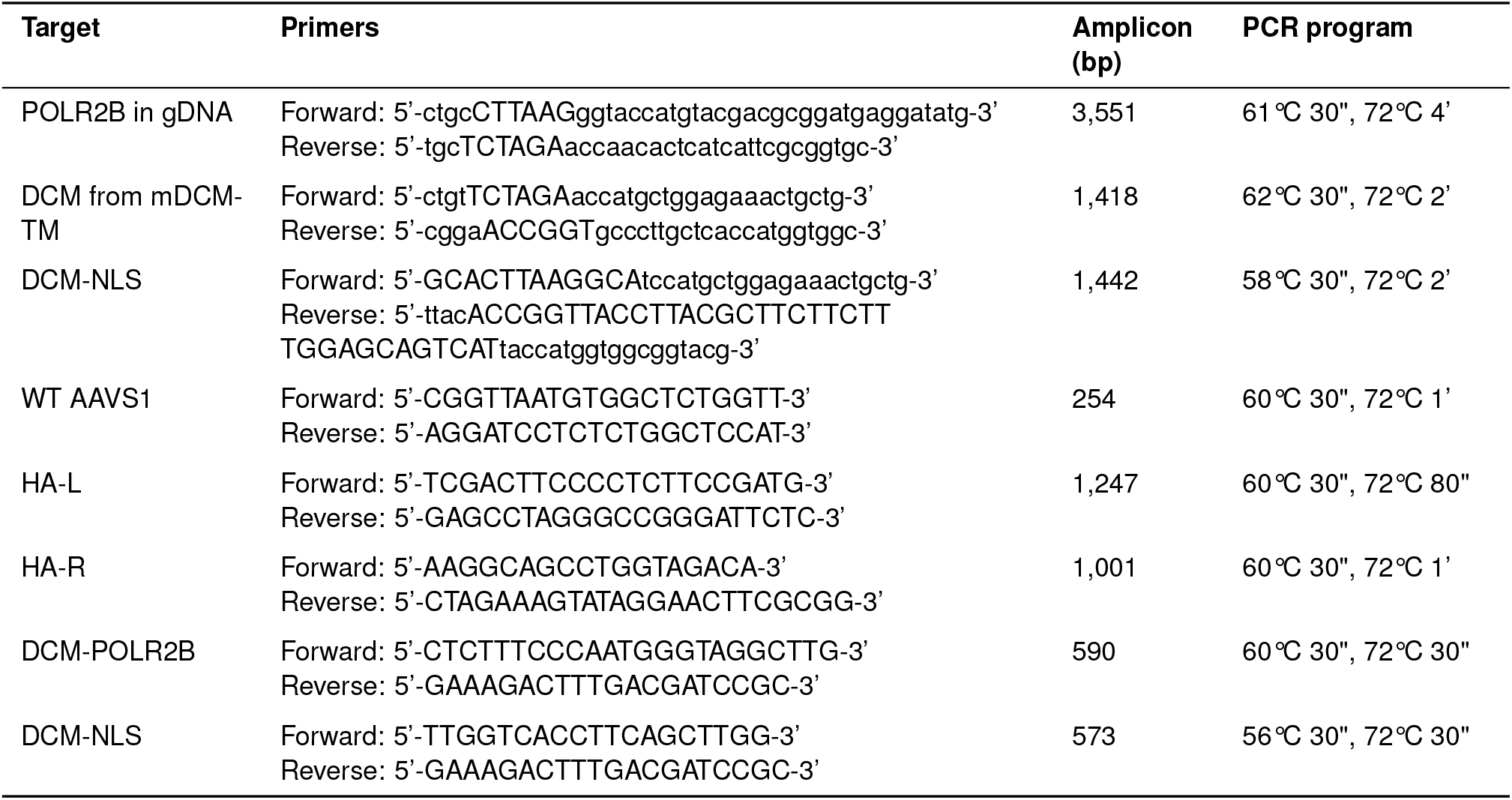
Primers used for PCR.

### Generation of human DCM-time machine cell lines

To generate the transgenic iPSC lines, the fusion genes were introduced in the genome using CRISPR/Cas9 and nucleofection. For nucleofection, the Human Stem Cell Nucleofector kit 2 (Lonza, VPH-5022) was used according to manufacturer’s instructions. Briefly, 1h before nucleofection medium was refreshed with culture medium supplemented with RevitaCell (Gibco, #A2644501). Cells were washed with D-PBS (Without Ca++ and Mg++; STEMCELL Technologies, 37350) and dissociated with StemPro Accutase Cell Dissociation Reagent (Gibco, a1110501), incubated for 7 min at 37 °C 5% CO2. To generate a single cell suspension, the cells were gently resuspended 5-10 times with a P1000.

Per nucleofection, 8 *×*10^5^ cells were collected and centrifuged 3 min at 1000 rpm. The pellet was washed with D-PBS, centrifuged 3 min 1000 rpm and D-PBS was completely removed. The cells were then resuspended in 82 µL of Human Stem Cell Nucleofector Solution 2 plus 18 µL of supplement, and this cell suspension was mixed with 2.5 µg of PX458-gRNA-hAAVS1 and 0.75 µg DCM-POLR2B donor vector (4:1). The PX458gRNA-hAAVS1 plasmid was generated by cloning the AAVS1sgRNA sequence (sgAAVS1_sense: 5’-CACCGGGGCCACTAGGGACAGGAT-3’, sgAAVS1_-antisense: 5’-AAACATCCTGTCCCTAGTGGCCCC-3’) into the pSpCas9(BB)-2A-Puro (PX458) plasmid (Addgene #48138). The mixture was transferred to an Amaxa cuvette and electroporated using the Amaxa 2b nucleofector with program B-016. Afterwards, using a Pasteur’s pipette, the cells were collected in 500 µL culture medium with RevitaCell and plated in a prewarmed coated 6 well dish with a final volume of 2 mL. After 24 hours, puromycin selection was started for 5 consecutive days with an increasing concentration of puromycin, 100 up to 200 ng/mL. When surviving colonies grew to an appropriate size, the colonies were picked for expansion and validation using genotyping, karyotyping and qRT-PCR. For genotyping, genomic DNA was isolated using the QuickExtract DNA extraction kit (Lucigen, QE0905) according to manufacturer’s instructions. Primer sequences are listed in Table 1.

### Validation of hDCM-TM in pluripotent iPSCs

Labeling of the hDCM-TM technology was validated with the WTC-DCM-*POLR2B* cell line using standard culture conditions for iPSC expansion. To induce the fusion gene the cells were treated with 4 µg/ml doxycycline hyclate (Sigma, D9891-25g) for 48 hours with daily medium changes. Induced cells were sorted based on a mCherry-high signal using a BD FACSAria II version 9.0.1. For MeD-seq, DNA was isolated with the Qiamp DNA micro kit (Qiagen, 56304) following manufacturer’s instructions. RNA for bulk RNA seq was isolated using the ReliaPrep RNA miniprep system (Promega, Z6012) following manufacturer’s instructions. Negative samples were not induced and not sorted.

To study DCM methylation propagation, WTC:DR cells were induced for 48 hours with doxycycline and subsequently labeled with the CellTrace Far Red Cell Proliferation kit (Invitrogen, C34571). Before staining, cells were dissociated in a single cell suspension using StemPro Accutase Cell Dissociation Reagent as described previously. Cells were collected in 1 ml PBS and incubated with 0.125 µM CellTrace solution for 15 minutes at 37 °C, protected from light. Afterwards, 5 ml of culture medium with Revitacell was added and cells were pelleted for 3 min at 1000 rpm. At this stage, cells were either plated in a Geltrex coated 6-well plate or collected for MeD-seq (*n* = 2) or analysis with a BD Fortessa flow cytometer (*n* = 3). The median far-red intensity was calculated with FlowJo 10.7.2 (*n* = 3). The collection procedure was repeated every 24 hours, for up to 72 hours. The DCM methylation propagation rate was calculated by relating the ratio of DCM and CpG reads to the number of cell divisions in multiple time intervals. DCM methylation propagation was calculated using the following equations:

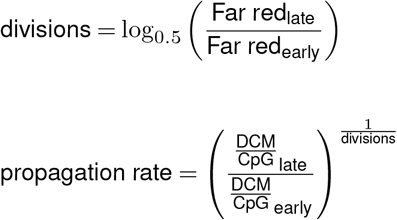

To predict how many cell divisions the DCM methylation labels remain detectable, we simulated the loss of DCM methylation-specific reads, as described previously [16]. Simulations were performed on iPSC clone 4 samples (*n* = 3) using a propagation rate of 72%. In brief, we calculated the fold change between in silico diluted and non-induced iPSC clone 4 samples (*n* = 3). We then determined the percentage of active genes with a fold change greater than 1, representing those that remain detectable after each division.

### Western blotting

Expression of the DCM-POLR2B fusion protein was visualized by Western blotting. +dox cells were induced for 48 hours with 4 µg/ml doxycycline hyclate (Sigma, D9891-25g) and collected using StemPro Accutase Cell Dissociation Reagent (Gibco, a1110501), as described previously. 1.1 *×*10^6^ mCherry positive cells were sorted on the FACSAriaIII (BD Biosciences). −dox samples were collected after Accutase treatment without FACS sorting. Cells were washed with PBS and total protein extraction was performed with RIPA buffer (Abcam, ab288006). Western blotting was performed as described previously with slight adjustments [16]. For DCM detection, 75 µg of total protein and for POLR2B detection 30 µg of total protein was loaded per lane. A Midi Format Transfer Pack with 0.2 µm PVDF membrane (Biorad #1704157) was used for blotting using a Trans-Blot Turbo Transfer System (Biorad #1704150) using a protocol for HIGH MW (2.5A, up to 25V for 10 min). The membrane was incubated for 30 minutes at RT in blocking buffer (1.3 g of non-fat dry milk in 50 ml of 1xTris-buffered saline) and probed with primary antibodies (1:1,000, DCM antibody: Cusabio CSB-PA365131XA01ENV, POLR2B Polyclonal Antibody: Thermo Fisher Scientific, PA5-30122) in blocking buffer + 0.1% Tween for 2 hours at room temperature. Then the membrane was incubated with anti-rabbit HRP secondary antibody (Sigma-Aldrich, A6154, 1:5,000) and a monoclonal anti-β-actin peroxidase antibody (1:10000, Sigma-Aldrich, A3854), as an internal control, in blocking buffer + 0.1% Tween for 1h at RT. Protein was visualized using the SuperSignal West Femto Maximum Sensitivity Substrate (Thermo Fisher Scientific, 34094) and an Amersham Imager 600 (Amersham Biosciences).

### Primitive gut tube differentiation

iPSCs were differentiated using the STEMdiff™ Pancreatic Progenitor Kit (STEMCELL Technologies, 05120) according to manufacturer’s instructions up until the primitive gut tube stage. To increase differentiation efficiency, the protocol was adapted by extending differentiation towards definitive endoderm with 24 hours, by medium refreshment. Differentiations were performed in a 12-well dish with 5 *×*10^5^ cells per differentiation.

The differentiation efficiency was established by flow cytometry using CXCR4 (PE anti-human CD184 (CXCR4) Antibody, Biolegend, #306505) and c-kit antibodies (APC anti-human CD117 (c-kit), Biolegend, #313205) on a BD LSRFortessa cell analyzer (BD Biosciences). Cell viability was determined using Hoechst 33342 (1:5000, Molecular Probes, H-3570). Analysis was performed in FlowJo 10.7.2. The differentiation protocol was further validated using real-time polymerase chain reaction (RT-qPCR). For this, cells were collected by incubation in rewarmed Accutase for 5 min at 37 °C 5% CO2 followed by RNA isolation using the ReliaPrep RNA miniprep system (Promega, Z6012) following manufacturer’s instructions. cDNA was prepared using GoTaq qPCR Master Mix (Promega, A6002). RT-qPCR was performed on a CFX384 real-time PCR-detection system (Bio-Rad). Values were normalized to hGAPDH, and fold changes in expression were calculated using the double delta Ct method. Primers can be found in Table 2. To apply the hDCM-Time machine, cells were induced for 24 hours with 4 µg/ml doxycycline hyclate at appropriate time points. Cells were collected using Accutase and DNA was isolated as described above.

**Table 2.**
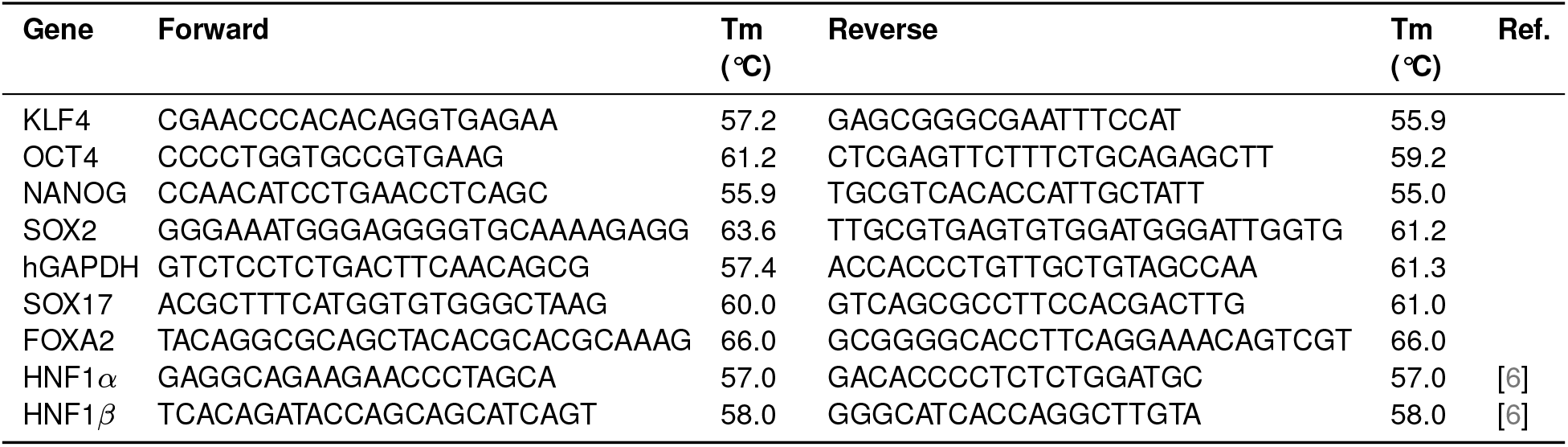
Primers used for gene expression analysis using qRT-PCR.

### Single-cell RNA sequencing

Viable single cells were FACS sorted into 384-well cell capture plates, from Single Cell Discoveries (Utrecht the Netherlands). Cell viability was determined using Hoechst 33342 (1:5000, Molecular Probes, H-3570). After sorting, plates were immediately spun and placed on dry ice. Plates were stored at −80°C. Single-cell RNA sequencing (scRNA-seq) was performed by Single Cell Discoveries according to an adapted version of the SORT-seq protocol [31]. The library was paired-end sequenced on an Illumina Nextseq™ 500, high output sequencer, using a 1×75 bp Illumina kit.

### scRNA-seq analysis

During sequencing, read 1 was assigned 26 base pairs and was used to identify the Illumina library barcode, cell barcode, and UMI. Read 2 was assigned 60 base pairs and used to map to the GRCh38 reference genome with STARSolo 2.7.10b [32]. Briefly, mapping and generation of count tables were automated using the STARSolo 2.7.10b aligner, soloFeatures set to ‘Gene’ and other settings set to default. Poisson correction was carried out on the count table matrix using a custom written Python script. This script converts the matrix (.mtx) to a dense format for element-wise processing. Afterward the script applies the Poisson correction based on count values.

All datasets were pre-processed separately using Scanpy v1.9.6 [33]. Low-quality cells were removed based on automatic thresholding using median absolute deviations (MAD), similar as described in Germain *et al*., 2020 [34]. Cells were flagged as outliers if they deviated more than 5 MADs in log(total_counts+1), log(n_-genes_by_counts+1) or pct_counts_in_top_20_genes, and were subsequently removed. For visualization, the datasets were merged, and a force-directed layout was generated using Harmony v0.1.5 [35]. First, counts were normalized, filtered for the top 5,000 highly variable genes and log-transformed using Harmony normalize_counts, hvg_genes and log_transform functions. An augmented affinity matrix was then created using time point information on affinities between iPSC and DE, and between DE and PGT (using the augmented_affinity_matrix function). Finally, this matrix was used to generate a force-directed layout for visualization using the force_directed_layout function.

Differentially expressed genes (DEGs) between cell types were identified using Scanpy. The dataset was prepared using Scanpy normalize_total, log1p, highly_-variable_genes, pca and neighbors functions with default settings. Differential expression analysis was then conducted using the rank_genes_groups function with Wilcoxon tests. DEGs were selected as genes with an adjusted *P* value < 0.01 and a log_2_(fold change) > 1. To compare expression levels between cell types, normalized counts per gene were extracted and averaged per cell type. These mean expression values were visualized in scatter plots comparing two cell types with all DEGs highlighted. Moreover, mean expression of all genes from each DEG group was plotted in a lineplot, showing changes in expression over time.

### MeD-seq

MeD-seq analysis was performed as previously described [17, 16]. At least 10 ng of DNA was digested by LpnPI (New England Biolabs). Sequencing libraries were prepped after LpnPI digestion using the ThruPLEX DNA-Seq HV kit (Takara, R400740). Using Pippin Prep system (Sage science) appropriate size fragments were selected. These libraries were subsequently sequenced on an Illumina NextSeq2000 sequencer, generating paired-end reads of 50 bases in length. For each condition, three biological replicates were sequenced.

### MeD-seq analysis

MeD-seq reads were filtered and categorized into DCM or CpG fastq files based on the presence of the LpnPI restriction site, located 13 to 17 base pairs from either the 5’ or 3’ end of the read. Reads from each fastq file were separately mapped to the GRCh38 reference genome using Bowtie2 v2.5.1 with default settings [36]. The number of reads overlapping each DCM or CpG site were counted using featureCounts v2.0.6 [37] with custom GTF files containing all DCM or CpG sites recognized by LpnPI, respectively. DCM sites were annotated using the annotatePeaks.pl script from Homer v4.11 [38]. The proportion of each annotation category per sample was determined by summing the read counts at each DCM site and assigning them to the corresponding annotation category.

### DCM methylation scores

To quantify DCM methylation levels per gene, which are corrected for differences in sequencing depth, propagation loss due to a chase period, and DCM induction level between samples, we calculated DCM methylation scores. First, we summed all DCM label counts at DCM sites overlapping the gene body of each gene in the Ensembl v108 annotation file. These counts were divided by the total number of DCM sites overlapping the gene, resulting in average number of DCM labels per DCM site. For each sample, these values were summed across all genes and divided by total number of MeD-seq reads to obtain a sample DCM level.

The average DCM level of the doxycycline-treated iPSC samples from clone 4 (*n* = 3) was set as the baseline level. For each doxycycline-induced sample, the fold change between its DCM level and the baseline level was calculated and used as a correction factor. For the non-induced samples, the correction factor was set at 1, indicating that the sample was corrected for sequencing depth but not for DCM methylation differences, which would lead to an overestimation of the background level. Finally, DCM scores for each gene were obtained by dividing the average number of DCM labels per DCM site by the correction factor.

### DCM profile plots

Profile plots were generated by dividing the genic region into three segments: upstream, gene body, and down-stream, with each segment further divided into 50 equal bins, totaling 150 bins. Each bin represents 1/50th of the total gene length. For each bin, DCM methylation scores from all DCM sites within that bin were extracted and averaged. The average profile was plotted using seaborn lineplot with bins on the x-axis and normalized counts on the y-axis, showing the mean as a solid line and the 95% confidence interval as shaded areas. Bins without DCM sites were excluded. The first bin was labeled as −100%, bin 50 as TSS, bin 100 as TES and the last bin as +100%.

### DCM labelling of genes

Differentially DCM methylated (DDM) genes were identified by comparing two conditions, iPSC^dox^ to DE^dox^, and DE^chase^ to PGT^chase^. For each gene, the DCM scores at sites overlapping its gene body were extracted and an independent *t* test was conducted between the two conditions. *P* values were then corrected using the Benjamini-Hochberg method. Genes were considered significantly DDM if they met the following thresholds: an adjusted *P* value < 0.05, and a log_2_(fold change) of < −0.5 (for downregulated genes) or > 0.5 (for upregulated genes).

DDM genes were validated using the scRNA-seq dataset, by calculating the mean expression of the list of genes across individual cells and plotting the results in the layout of the scRNA-seq data. These mean expression values were further summarized in violin plots to show differences in expression between cell types. Statistical comparisons between cell types were performed using Mann-Whitney tests, and *P* values were corrected for multiple testing using the Benjamini-Hochberg method.

### DCM labelling of enhancers

Enhancers were called from the DCM-TM dataset by performing a differential analysis between iPSC^dox^ and DE^dox^. First, intergenic DCM sites located at least 1 kb away from gene bodies were selected as potential enhancer sites. Sliding windows were constructed consisting of adjacent DCM sites. For each window, we counted the number of DCM labels and summed these values. To identify significantly different windows, a DESeq2 v1.42.0 analysis [39] was performed comparing iPSC^dox^ and DE^dox^ samples. Dispersions were estimated using a parametric fit and fold changes were shrunk using the ashr v2.2-63 method [40]. Significant windows were defined as windows with an adjusted *P* value < 0.05 and a log_2_(fold change) < −1 or > 1. Overlapping windows were then merged to obtain final enhancer regions.

DCM labelling of enhancer regions in the chase samples was validated by plotting the DCM scores of all significant regions across the dox (all replicates) and chase (mean of *n* = 3) samples. Moreover, boxplots displaying DCM scores at iPSC- and DE-specific enhancers were generated for each condition (mean of *n* = 3). Statistical comparisons between conditions were performed using Mann-Whitney tests, and *P* values were corrected for multiple testing using the Benjamini-Hochberg method.

### CpG methylation analysis

Differentially CpG-methylated regions (DMRs) were identified across four genomic regions: (1) CpG islands downloaded from UCSC, (2) promoter regions defined as TSS ±1kb, (3) gene body regions defined as TSS + 1kb to TES, and (4) DCM enhancer regions. The analysis included all genes from Ensembl v108, except those smaller than 2kb and regions on unknown chromosomes. To quantify CpG methylation, a SAF file was generated containing CpG sites recognized by LpnPI that overlapped the regions of interest. For each sample, CpG read counts per region were obtained using featureCounts v2.0.6. These counts were analyzed in DESeq2 v1.42.0 using the parametric fitting approach. Differential methylation was assessed between iPSCs and DE cells and between DE and PGT cells using the results function, after which the fold changes were shrunken using lfcShrink with the ashr v2.2-63 approach. Significant DMRs were defined as those with an adjusted *P* value < 0.05 and log_2_(fold change) < −0.25 or > 0.25.

To assess the functional relevance of DMRs, all promoter DMRs were extracted, and the expression levels of their corresponding genes were analyzed in the scRNA-seq dataset. Gene set enrichment was calculated using score_genes function from Scanpy, and the enrichment scores per cell were visualized in the scRNA-seq layout. These scores were further summarized per cell type in a violin plot. Additionally, two marker genes with significant promoter DMRs, *HNF4A* and *OCT6*, were examined in detail. Their normalized scRNA-seq expression levels per cell type and promoter methylation scores per sample were plotted. CpG methylation scores were normalized using size factors calculated by DESeq2. Similarly, enhancer DMRs were further examined by selecting DCM enhancers with significant CpG methylation differences. These enhancers were categorized by cell type-specify in the DCM data. For each category, DCM methylation levels were plotted, along with CpG methylation levels, normalized using DESeq2-derived size factors. Statistical comparisons between cell types were performed using Mann-Whitney tests, with *P* values corrected for multiple testing using the Benjamini-Hochberg method.

### Genome browser views

Genome browser views were created using IGV v2.18.2 [41]. Tracks with DCM scores per sample were generated using deepTools v3.5.5 [42] bamCoverage using a bin size of 1 and a scale factor of 1/correction factor (see “DCM methylation scores”). Tracks with CpG methylation were also created using bamCoverage, with the CpG read count per million as scale factor. Average tracks per condition were generated by summing the scores per site across replicates using UCSC bigWigmerge v2 [43] and then dividing the sum by the number of replicates with a custom Python script.

### Bulk RNA sequencing

Total RNA was extracted from doxycycline-treated and -untreated iPSCs (*n* = 3). Sequencing libraries were prepared using TruSeq stranded mRNA library preparation method from Illumina. These libraries were subsequently sequenced on an Illumina NextSeq2000 sequencer. Paired-end clusters were generated of 50 bases in length, resulting in 40 million reads per sample.

### Bulk RNA-seq analysis

RNA-seq reads were trimmed and quality-filtered using Trim Galore v0.6.10. These processed reads were then mapped to the GRCh38 reference genome using HISAT2 v2.2.1 with default settings [44]. Transcript counts were obtained with featureCounts v2.0.6 (-s 2 -t exon), utilizing the Ensembl v108 annotation file. Differential expression analysis between the doxycycline-treated and -untreated iPSCs was performed using DESeq2 v1.42.0, employing a parametric fit for dispersion estimation. Calculated fold changes were shrunk using the ashr v2.2-63 method. Differentially expressed genes were selected based on an adjusted *P* value < 0.05 and a log_2_(fold change) < −1 or > 1. The results were further filtered to retain only characterized genes (i.e. those with annotated names, excluding Gm and ribosomal genes) and visualized in an MA plot, with the log_10_ of the base mean on the x-axis and the log_2_(fold change) on the y-axis.

For comparison with the MeD-seq data from iPSCs, transcripts per million (TPM) were calculated for each gene in each sample. For both the doxycycline-treated and -untreated samples, scatter plots were created showing the average RNA-seq TPM and average MeD-seq normalized counts. The correlation between the two datasets was assessed using Pearson correlation coefficients. Furthermore, based on the average expression levels across all RNA-seq samples (*n* = 6), genes were categorized in three equal expression groups: low, medium and high. Profile plots showing the DCM labelling along the gene body were generated for each of these three categories separately. For visualization, bigwig files were generated using deepTools v3.5.5 bamCoverage with a bin size of 1 and CPM as normalization method.

### Correlations between iPSC, DE and PGT datasets

Spearman correlation coefficients were calculated between MeD-seq, RNA-seq and scRNA-seq samples. To focus on cell type-specific differences, we selected DEGs from the scRNA-seq analysis and computed correlation coefficients using only these genes. DCM scores per gene from the MeD-seq samples were extracted and averaged across conditions (*n* = 3). In the RNA-seq analysis, mean TPM values were calculated for the C4 clone under both uninduced and induced conditions (*n* = 3). For scRNA-seq, mean expression values for each cell type were calculated using a pseudobulk approach. Spearman correlation coefficients were then calculated between these values. Additionally, correlations between the MeD-seq samples and each cell in the scRNA-seq dataset were calculated and visualized in the layout of the scRNA-seq data.

### ChIP-seq analysis

ChIP-seq data from iPSC, DE and PGT samples were re-analyzed to assess histone modification and RNA polymerase II (RNAP II) enrichment at DCM-labelled sites and genes. For iPSCs, datasets for H3K4me3, H3K27ac, H3K27me3 H3K36me3, and RNAP II were obtained from GSE220103, GSE158382, and GSE145964 [45, 46, 18]. For DE and PGT, data for H3K4me3, H3K27ac, and H3K27me3 were retrieved from GSE149148 [19]. Reads were downloaded from the SRA database and trimmed using Trim Galore v0.6.10. The processed reads were then mapped to the GRCh38 reference genome using Bowtie2 v2.5.1 with default settings. Peak calling was performed using MACS3 v3.0.1 callpeak [47] with default parameters for all analyses, except for H3K36me3 and H3K27me3, where broad peaks were called using the –broad option. Additionally, the -f BAMPE setting was applied for all paired-end datasets. Tracks for plotting were generated using MACS3 bdgcmp with the ppois method.

Detected DCM labels were validated using ChIP-seq datasets. A profile plot of POLR2B enrichment along the gene body was generated with deepTools v3.5.5 using the computeMatrix (scale-regions mode, averageTypeBins mean, 50bp bins, 2kb up-/downstream) and plotProfile functions. To assess enhancer labeling, DCM scores at H3K27ac peaks in iPSCs and DE cells were plotted. For these plots, computeMatrix was applied in reference-point mode (500bp bins, averageTypeBins mean, 5kb up-/downstream). Missing values in bins were replaced with 0 to ensure visualization of regions without DCM methylation labels. The resulting heatmap was generated using deepTools plotHeatmap, with regions sorted based on DCM enrichment to highlight peaks with and without labeling.

### Motif analysis

Motif analyses were performed on the enhancer regions specific to iPSCs and DE cells, identified from the DCM enhancer analysis. To account for the recognition sequences of LpnPI, a motif file was created containing sequences that should be masked during the analysis. These sequences (CCG, CGG, GCGC, CCAGG, and CCTGG) were processed using the seq2profile.pl script from Homer v4.11. Subsequently, the findMotifsGenome.pl script from Homer was used with the “-size given” option and the motif file to mask these sequences. The results were sorted by *P* value, and the top 8 motifs with the highest significance in iPSCs and DE cells were visualized using bar plots. Additionally, motif analyses were performed on H3K27ac peaks in iPSC and DE samples using the findMotifsGenome.pl script with the “-size given” option. Significant motifs (*P* value < 0.05) from this analysis were selected and compared with the top motifs identified in the DCM enhancer analysis for the corresponding cell type.

## Resource availability

### Lead contact

Further information and requests for resources and reagents should be directed to and will be fulfilled by the lead contact, Joost Gribnau.

### Materials availability

Generated cell lines used in this study can be made available.

### Data availability

All raw and processed high-throughput sequencing data (MeD-seq, bulk RNA-seq and scRNA-seq) generated in this study have been submitted to the NCBI Gene Expression Omnibus (GEO) under accession number GSE304237. Moreover, ChiP-seq datasets from GSE158382, GSE220103, GSE145964 and GSE149148 were reanalyzed. The code used for data analysis is available upon reasonable request.

### Code availability

The code used for data analysis is generally available upon reasonable request, excluding scripts for processing raw MeD-seq data, which are restricted due to licensing agreements between Erasmus Medical Center and commercial partners.

## Acknowledgements

We thank the members of the Erasmus MC Developmental Biology department for valuable discussions, and Menno Creyghton for critically reading the manuscript. J.G. and M.E.v.L. were supported by an NWO Psider grant (nr. 40-46800-98-015-10250022120002).

## Author contributions

M.E.v.L., B.F.T. and J.G. conceived the study, performed the experiments, analyzed the data, and wrote the manuscript. E.W. and E.S. assisted in FACS analysis. R.B. performed the MeDseq data preprocessing. C.G. aided in the generation of transgenic iPSC lines.

## Supplementary Information

**Figure S1.**
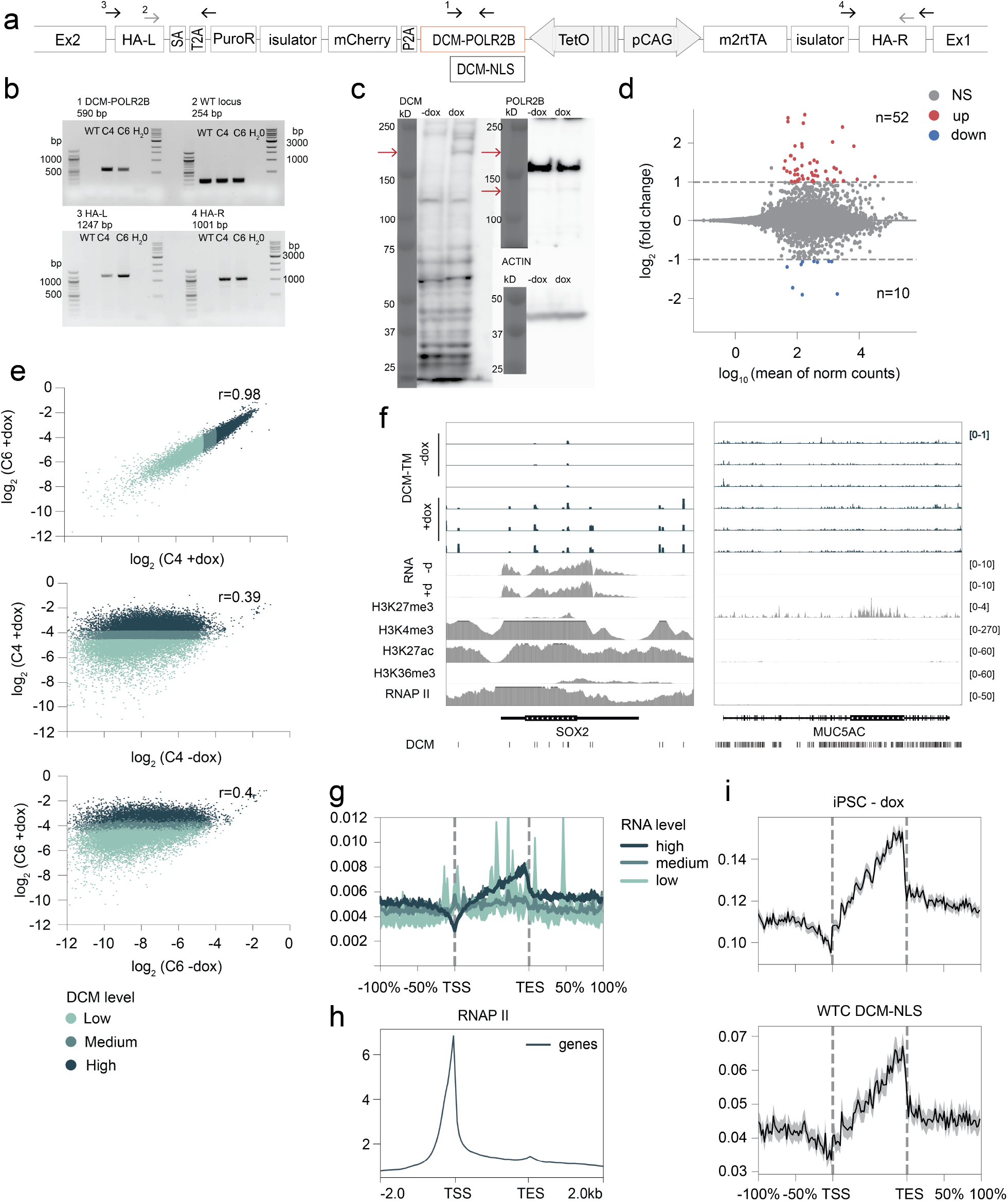
**a**. Introduction of DCM-*POLR2B* and DCM-NLS in the *AAVSI* locus using CRISPR/Cas9. Arrows indicate the location of primers used for genotyping. HA-L, left homology arm; HA-R, right homology arm; Ex, exon; SA, Splice acceptor site; PuroR, Puromycin Resistance cassette. **b**. Genotyping by PCR of DCM-*POLR2B* integration in *AAVS1* locus. Primer pairs and PCR programs are provided in **a** and Table 1. WT, wild type band. **c**. Western blot of DCM-POLR2B iPSCs with and without doxycycline (dox) induction, DCM-POLR2B-mCherry 213.3 kDa, DCM-POLR2B 186.6 kDa, POLR2B 134.3 kDa, actin 42 kDa. **d**. Differential expression analysis between RNA-seq data of −dox and +dox DCM-*POLR2B* iPSCs. Differentially expressed genes (DEGs) are highlighted in red and blue, indicating up- and downregulated genes, respectively. NS, not significant. **e**. Scatter plots showing correlation between average DCM scores (of *n* = 3) for two DCM-*POLR2B* iPSC cell lines (top), −dox and +dox of clone 4 (center), and −dox and +dox of clone 6 (bottom). Genes are highlighted according to three DCM score groups in the C4 +dox samples, indicating low, medium and high levels of labelling. Pearson correlation coefficient is indicated. **f**. Genome browser view of DCM, RNA-seq and ChIP-seq tracks at loci of pluripotency gene *SOX2* and the inactive gene *MUC5AC*. d, doxycycline. RNAP II: RNA polymerase II. **g**. Profile plots of DCM methylation labelling in DCM-*POLR2B* −dox iPSCs, showing distribution of DCM reads along the gene body, upstream of the transcription start site (TSS, −100% of the gene length), and downstream of the transcription end site (TES, +100%). Genes are split in categories based on gene expression levels in bulk RNA-seq data. Mean DCM scores with 95% confidence intervals are shown. **h**. Profile plot of POLR2B ChIP-seq enrichment at all genes, showing 2kb upstream of the TSS to 2kb downstream of the TES. **i**. Profile plots of DCM methylation labeling in DCM-*POLR2B* −dox iPSCs (top) and DCM-NLS +dox iPSCs (bottom). Mean DCM scores with 95% confidence intervals are shown.

**Figure S2.**
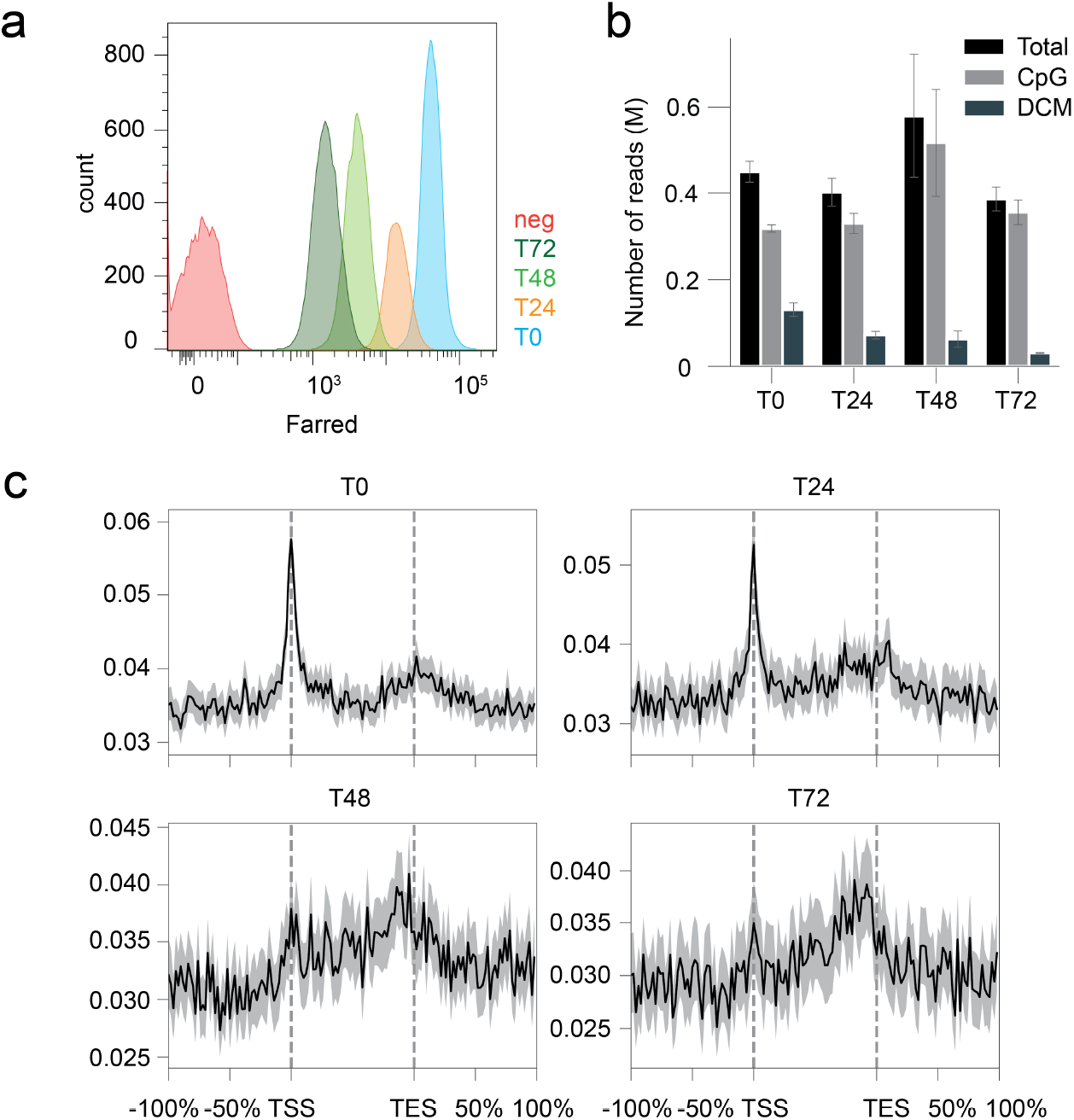
**a**. CellTrace intensity up to 72 hours after stain with label. T, timepoint. **b**. Number of sequenced MeD-seq reads, including total, CpG-specific and DCM-specific reads in the propagation pulse-chase experiment. Mean of n=2 with 95% confidence interval is shown in millions for each timepoint. **c**. Profile plots of DCM methylation labelling across all genes at each time point of the propagation pulse-chase experiment. Mean DCM scores of n=2 with 95% confidence intervals are shown.

**Figure S3.**
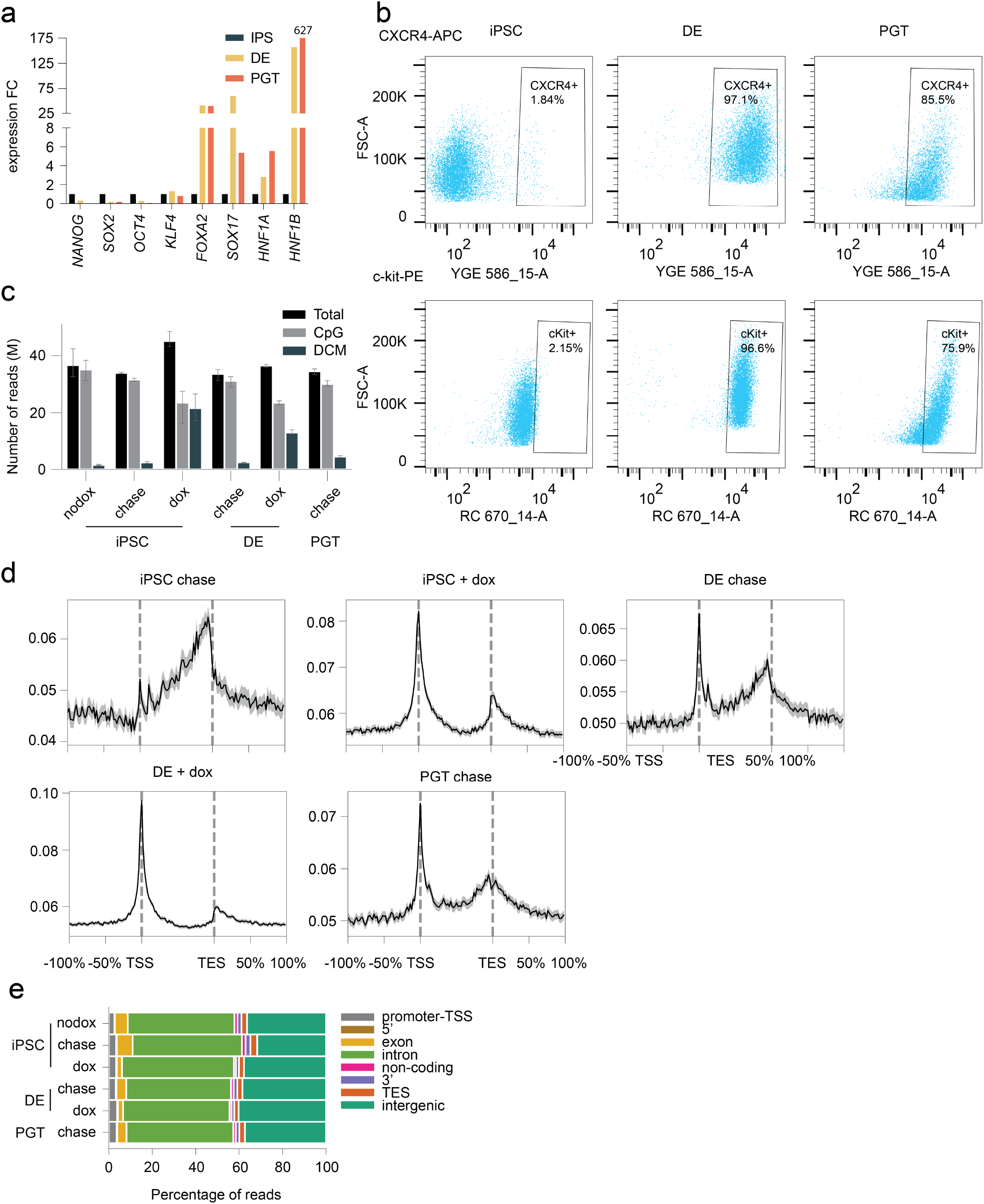
**a**. RT-qPCR analysis of marker genes during PGT differentiation. Fold change (FC) relative to iPSC expression is shown. **b**. Flow cytometry analysis of DE marker genes CXCR4 (top) and cKIT (bottom) in iPSCs (left), DE (center), and PGT cells (right). **c**. Number of sequenced MeD-seq reads, including total, CpG-specific and DCM-specific reads in the PGT differentiation experiment shown in Figure 3a. Mean of *n* = 3 with 95% confidence interval is shown in millions. **d**. Profile plots of DCM methylation labelling in the PGT differentiation experiment, showing distribution of DCM reads along the gene body, upstream of the transcription start site (TSS, −100% of the gene length), and downstream of the transcription end site (TES, +100%). Mean DCM scores with 95% confidence intervals are shown. **e**. Genomic annotation of DCM-specific reads in the PGT differentiation samples.

**Figure S4.**
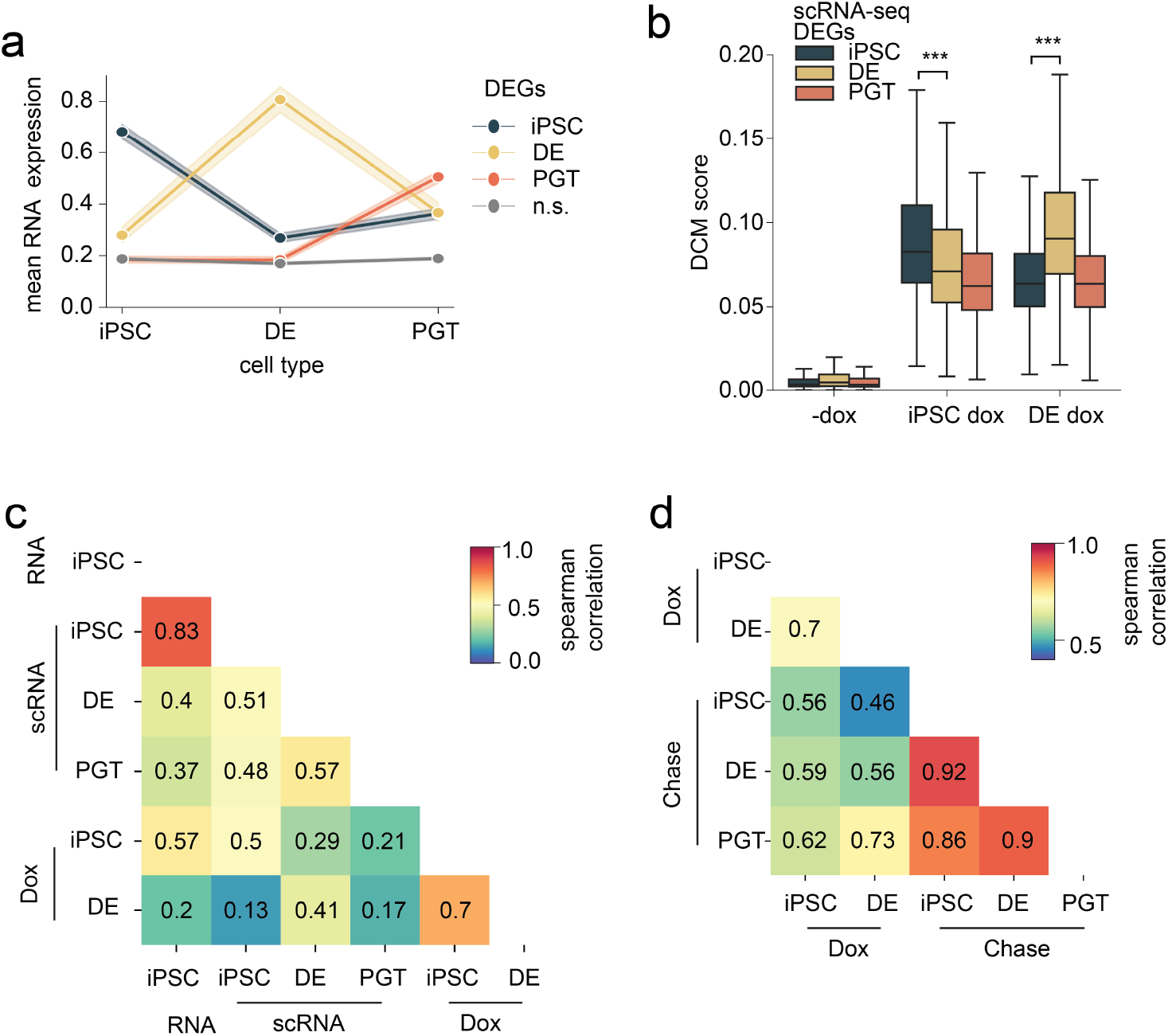
**a**. Mean pseudobulk scRNA-seq expression levels of DEGs between cell types (from Figure 3d) and all non-significant (n.s.) genes. Mean expression with 95% confidence interval is shown. **b**. DCM scores of scRNA-seq DEGs (from Figure 3d) in −dox and +dox iPSCs. The box plots display the median (black line), the interquartile range (box limits) and 1.5x of the interquartile range (whiskers). *P* values were calculated using Mann-Whitney tests and corrected for multiple testing using the Benjamini-Hochberg approach (*** *P* value < 0.001). **c**. Heatmap showing Spearman correlation between RNA-seq, scRNA-seq pseudobulk, and DCM dox samples, calculated using only the scRNA-seq DEGs shown in Figure 3d. **d**.Heatmap showing Spearman correlation between DCM scores of the dox and chase samples, calculated using only the scRNA-seq DEGs of Figure 3d.

**Figure S5.**
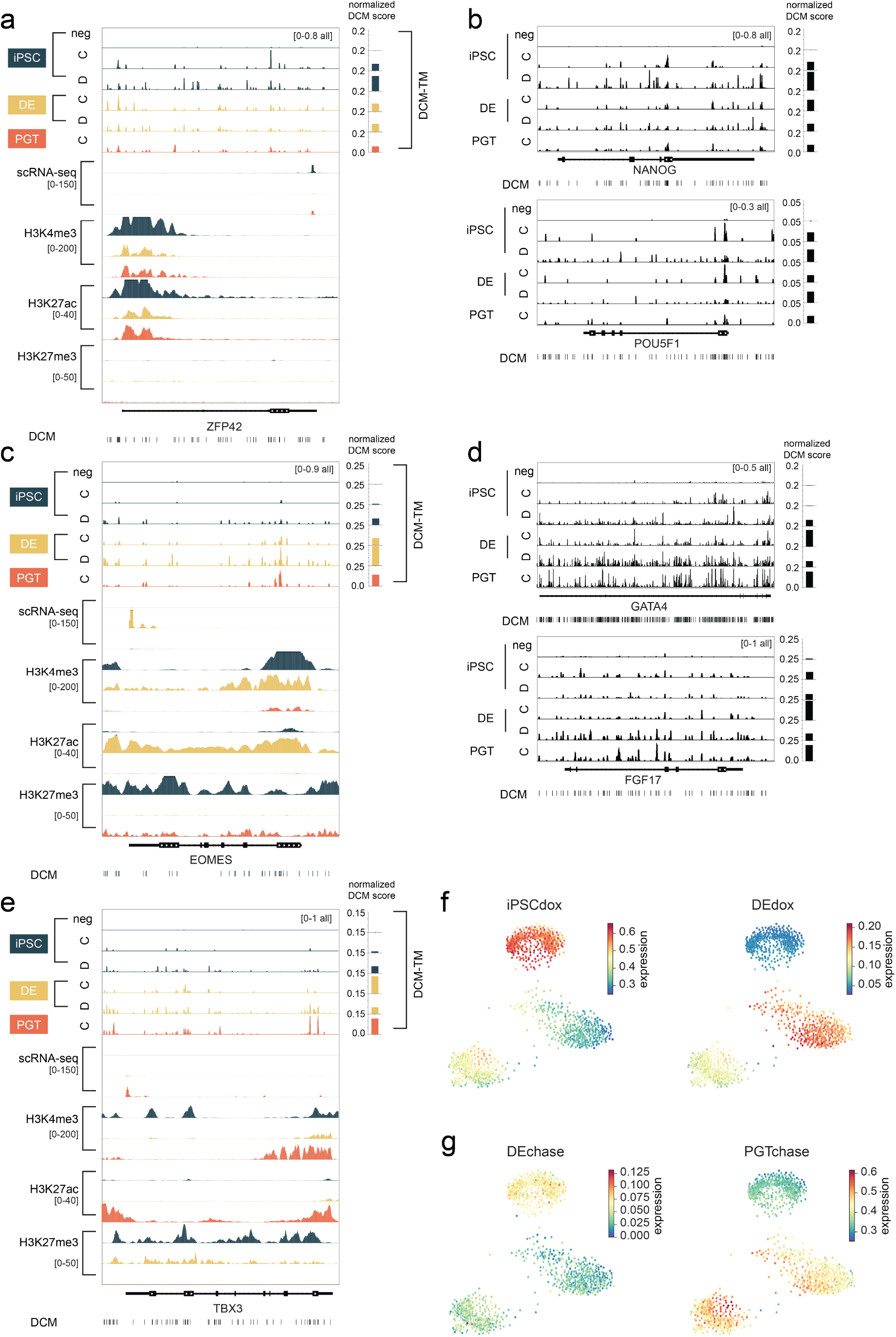
**a**,**c**,**e**. Genome browser view of iPSC marker *ZFP42* (**a**), DE marker *EOMES* (**c**), and PGT marker *TBX3* (**e**). Tracks show DCM methylation (average of *n* = 3), scRNA-seq pseudobulk tracks of iPSC, DE an PGT cells, and H3K4me3, H3K27ac and H3K27me3 enrichment iPSC, DE and PGT cells. The barplots on the right quantify all DCM scores overlapping each gene per sample. C, chase; D, dox. **b**,**d**. Genome browser view of DCM methylation labelling at loci of pluripotency markers (*NANOG* and *POU5F1*, **b**) and DE markers (*GATA4* and *FGF17*, **d**). The barplots on the right quantify all DCM scores overlapping each gene per sample. C, chase; D, dox. **f**,**g**. Average expression of differentially DCM methylated genes (DDMs) between iPSC^dox^ and DE^dox^ (**f**) and between between DE^chase^ and PGT^chase^ (**g**) on scRNA-seq layout.

**Figure S6.**
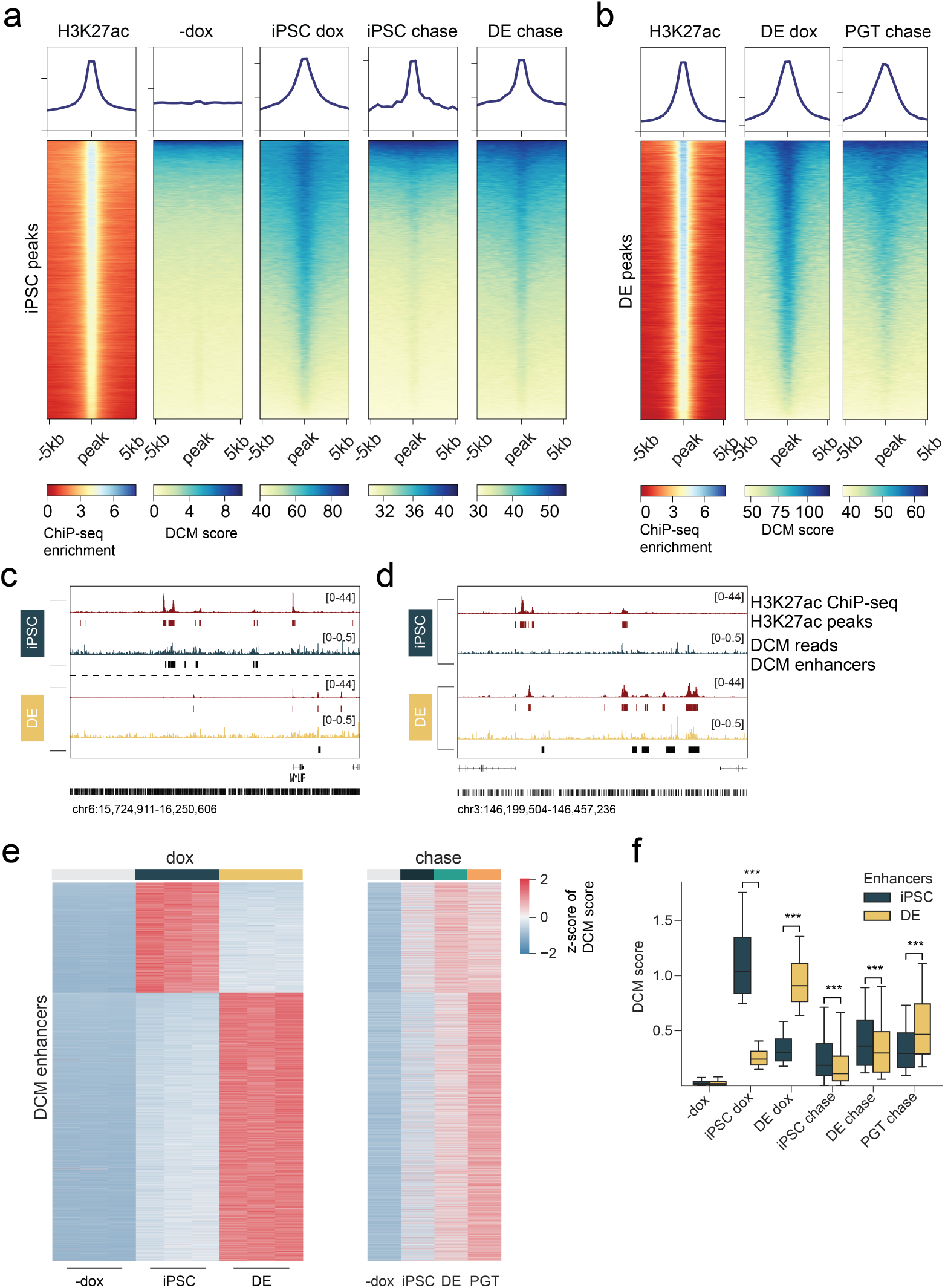
**a**. Heatmap showing H3K27ac enrichment in iPSCs and DCM labels of all related MeD-seq samples at H3K27ac peaks in iPSCs. Profile plots have the same y-axis range as the corresponding heatmap. DCM scale *×*10^3^. **b**. Heatmap showing H3K27ac enrichment in DE cells and DCM labels of all related MeD-seq samples at H3K27ac peaks in DE cells. Profile plots have the same y-axis range as the corresponding heatmap. DCM scale *×*10^3^. **c-d**. Genome browser view of iPSC enhancers in an intergenic region on chromosome 6 (**c**) and DE enhancers in an intergenic region on chromosome 3 (**d**). Tracks shown correspond to those in Figure 5a. **e**. Heatmap of DCM methylation of enhancers in dox (left) and chase (right) samples, showing z-scores of DCM scores (dox) and z-scores of average DCM scores (chase, *n* = 3). **f**. Quantification of **e**, showing DCM scores at iPSC enhancer regions (blue) and DE enhancer regions (yellow). The box plots display the median (black line), the interquartile range (box limits) and 1.5x of the interquartile range (whiskers). *P* values were calculated using Mann-Whitney tests and corrected for multiple testing using the Benjamini-Hochberg method (*** p<0.001).

**Figure S7.**
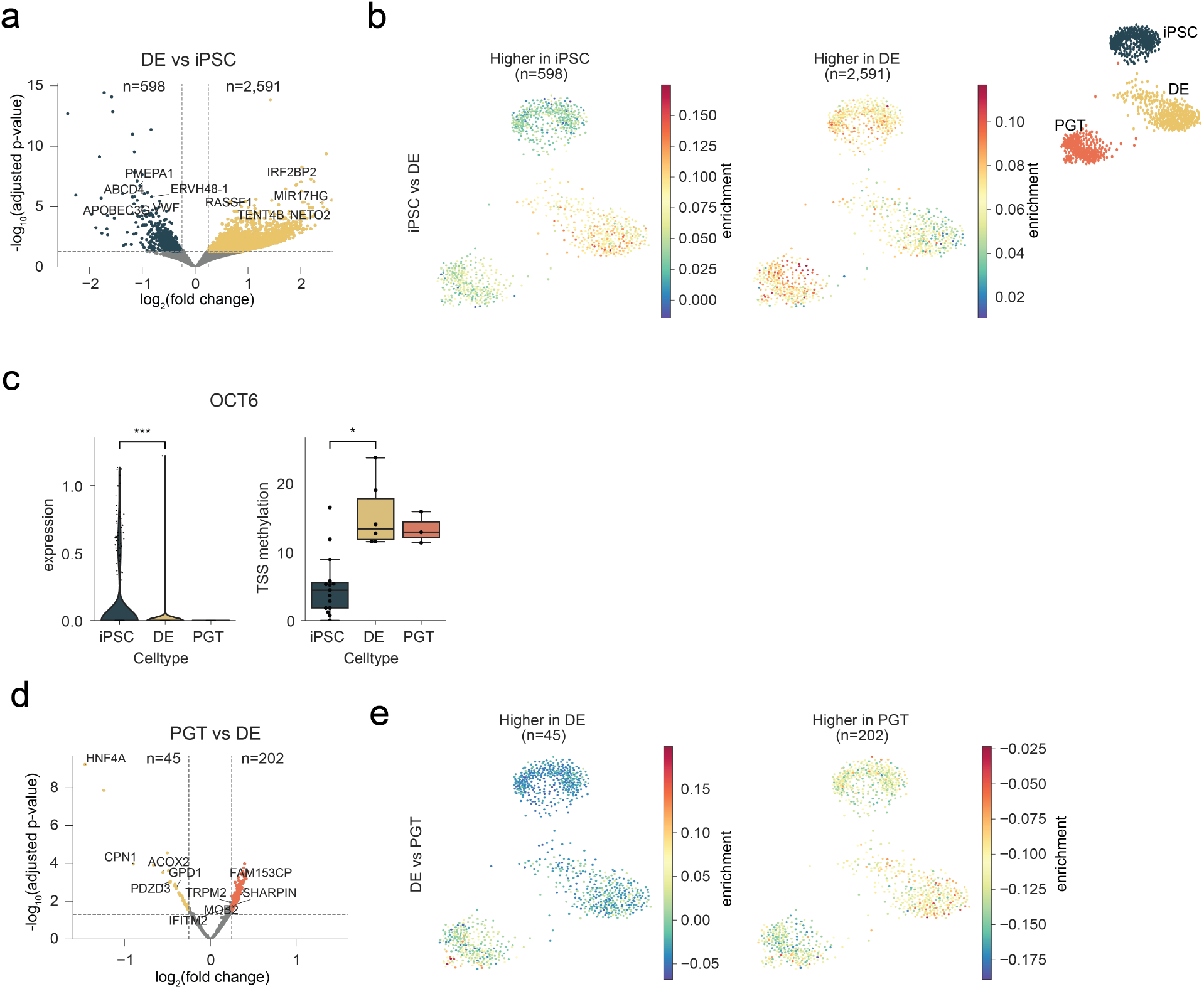
**a**. Volcano plot of differentially CpG-methylated promoter regions between iPSCs and DE cells. **b**. Enrichment of genes with differentially methylated promoters between iPSCs and DE cells (**a**) visualized on the scRNA-seq layout. **c**. RNA expression of *OCT6* (left) and CpG methylation at the promoter (right). *P* values were calculated using Mann-Whitney tests (* *P* value < 0.05, ** *P* value < 0.01, *** *P* value < 0.001). **d**. Volcano plot of differentially CpG-methylated promoter regions between DE and PGT cells. **e**. Enrichment of genes with differentially methylated promoters between DE and PGT cells (**d**) visualized on the scRNA-seq layout.

## Supplementary Tables

**Table S1.** Statistics of MeD-seq samples.

| Sample | Total reads | DCM reads | CpG reads | Correction factor | DCM/CpG |
| --- | --- | --- | --- | --- | --- |
| iPSCs<sup>-dox</sup> C4 1 | 42,414,674 | 1,756,919 | 40,657,755 | 42.41 | 0.04 |
| iPSCs<sup>-dox</sup> C4 2 | 32,645,592 | 1,431,276 | 31,214,316 | 32.65 | 0.05 |
| iPSCs<sup>-dox</sup> C4 3 | 34,929,672 | 1,383,492 | 33,546,180 | 34.93 | 0.04 |
| iPSCs<sup>dox</sup> C4 1 | 42,715,841 | 26,629,059 | 16,086,782 | 55.31 | 1.66 |
| iPSCs<sup>dox</sup> C4 2 | 48,475,994 | 20,953,967 | 27,522,027 | 43.32 | 0.76 |
| iPSCs<sup>dox</sup> C4 3 | 44,165,595 | 17,234,310 | 26,931,285 | 35.84 | 0.64 |
| iPSCs<sup>-dox</sup> C6 1 | 36,506,398 | 1,591,859 | 34,914,539 | 36.51 | 0.05 |
| iPSCs<sup>-dox</sup> C6 2 | 34,477,290 | 1,273,347 | 33,203,943 | 34.48 | 0.04 |
| iPSCs<sup>-dox</sup> C6 3 | 34,594,914 | 1,483,964 | 33,110,950 | 34.59 | 0.04 |
| iPSCs<sup>dox</sup> C6 1 | 38,526,918 | 16,985,836 | 21,541,082 | 36.50 | 0.79 |
| iPSCs<sup>dox</sup> C6 2 | 38,250,331 | 8,978,049 | 29,272,282 | 18.78 | 0.31 |
| iPSCs<sup>dox</sup> C6 3 | 41,932,304 | 10,920,096 | 31,012,208 | 22.66 | 0.35 |
| iPSCs DCM-NLS | 32,814,027 | 1,698,705 | 31,115,322 | 4.59 | 0.05 |
| iPSCs CellTrace 0h 1 | 473,045 | 146,485 | 326,560 | 0.52 | 0.45 |
| iPSCs CellTrace 0h 2 | 424,895 | 115,254 | 309,641 | 0.38 | 0.37 |
| iPSCs CellTrace 24h 1 | 370,062 | 63,320 | 306,742 | 0.22 | 0.21 |
| iPSCs CellTrace 24h 2 | 433,831 | 80,944 | 352,887 | 0.31 | 0.23 |
| iPSCs CellTrace 48h 1 | 722,456 | 81,646 | 640,810 | 0.31 | 0.13 |
| iPSCs CellTrace 48h 2 | 436,528 | 44,152 | 392,376 | 0.17 | 0.11 |
| iPSCs CellTrace 72h 1 | 413,751 | 29,769 | 383,982 | 0.12 | 0.08 |
| iPSCs CellTrace 72h 2 | 358,726 | 31,817 | 326,909 | 0.11 | 0.10 |
| iPSCs<sup>chase</sup> 1 | 34,190,998 | 2,079,124 | 32,111,874 | 5.43 | 0.06 |
| iPSCs<sup>chase</sup> 2 | 33,699,370 | 2,686,999 | 31,012,371 | 7.16 | 0.09 |
| iPSCs<sup>chase</sup> 3 | 33,967,760 | 2,335,542 | 31,632,218 | 5.99 | 0.07 |
| DE<sup>chase</sup> 1 | 31,324,302 | 2,308,304 | 29,015,998 | 5.31 | 0.08 |
| DE<sup>chase</sup> 2 | 35,128,406 | 2,490,389 | 32,638,017 | 5.73 | 0.08 |
| DE<sup>chase</sup> 3 | 34,332,371 | 2,432,121 | 31,900,250 | 5.48 | 0.08 |
| DE<sup>dox</sup> 1 | 36,910,216 | 14,051,164 | 22,859,052 | 30.03 | 0.61 |
| DE<sup>dox</sup> 2 | 35,966,488 | 12,596,433 | 23,370,055 | 26.74 | 0.54 |
| DE<sup>dox</sup> 3 | 36,657,466 | 12,481,752 | 24,175,714 | 26.33 | 0.52 |
| PGT<sup>chase</sup> 1 | 33,464,132 | 4,124,227 | 29,339,905 | 9.68 | 0.14 |
| PGT<sup>chase</sup> 2 | 36,014,703 | 4,792,058 | 31,222,645 | 11.09 | 0.15 |
| PGT<sup>chase</sup> 3 | 34,056,712 | 4,389,016 | 29,667,696 | 10.13 | 0.15 |

**Table S2.** Statistics of bulk RNA-seq samples.

| Sample | Total reads | Mapped reads (%) |
| --- | --- | --- |
| iPSCs<sup>-dox</sup> 1 | 53,453,806 | 98.83 |
| iPSCs<sup>-dox</sup> 2 | 52,223,120 | 98.90 |
| iPSCs<sup>-dox</sup> 3 | 39,963,748 | 98.93 |
| iPSCs<sup>dox</sup> 1 | 45,088,918 | 98.77 |
| iPSCs<sup>dox</sup> 2 | 38,776,834 | 98.36 |
| iPSCs<sup>dox</sup> 3 | 42,903,044 | 98.15 |

**Table S3.** DEGs between cell types in scRNA-seq dataset. The 50 most significant DEGs per cell type are shown.

| iPSC | DE | PGT |
| --- | --- | --- |
| SFRP2 | FGF17 | TTR |
| ERVH-1 | PTGR1 | FN1 |
| DNMT3B | LEFTY1 | HMGCS2 |
| TERF1 | LAPTM4B | NPY |
| SOX2 | EOMES | APOA2 |
| DPPA4 | TMSB10 | IGFBP3 |
| ESRG | CNTNAP2 | CST3 |
| UTF1 | LEFTY2 | PCAT14 |
| PSAT1 | PLSCR2 | LINC00261 |
| DANCR | LINC01356 | TBX3 |
| SEPHS1 | CER1 | ID2 |
| NANOG | SERPINB9 | FLRT3 |
| HSPE1 | CYP26A1 | PRTG |
| NPM1 | SOX17 | CKB |
| FABP5 | BEX3 | VCAN |
| C1QBP | RPL10 | PTPN13 |
| DDX21 | SFRP1 | FZD4 |
| CD9 | APLP2 | LAMB1 |
| PHC1 | PKM | ID1 |
| PDPN | RPLP2 | GATA3 |
| HSPD1 | RPS19 | COL4A6 |
| VASH2 | KCNK12 | HLA-B |
| CYCS | LINC00458 | CADM1 |
| UGP2 | MIXL1 | IMPAD1 |
| H2AFZ | RPL28 | DLK1 |
| POU5F1 | OTX2 | ADD3 |
| DPYSL2 | MDK | ID3 |
| ZSCAN10 | KRT8 | TTC3 |
| RAN | CCKBR | RGS2 |
| PAICS | SREK1IP1 | S100A10 |
| CNMD | RHOBTB3 | KRT19 |
| POLR3G | MTRNR2L1 | MCC |
| JARID2 | IFITM3 | IDH2 |
| THY1 | GATA6 | BAMBI |
| PTPRZ1 | ARL4D | S100A11 |
| FOXD3-AS1 | GSTP1 | SYTL5 |
| CCT5 | CAVIN3 | GSTA2 |
| PGAM1 | AC068587.4 | COL9A2 |
| RBPMS | SERHL2 | ZFP36L2 |
| B3GNT7 | SPARC | TSHZ2 |
| TRIM28 | IGFBPL1 | VIL1 |
| TUBB3 | TRDN | CYP4X1 |
| SET | LY6E | ALCAM |
| SRSF2 | MYL6 | AMOT |
| YBX1 | TNFSF14 | CXXC4 |
| HSP90AB1 | LHX1 | CNKSR2 |
| CCT6A | NXN | COL4A5 |
| RCC2 | APOC1 | CGN |
| LINC00678 | DUSP4 | NR2F2 |
| PCNA | MGST2 | APELA |

**Table S4.**
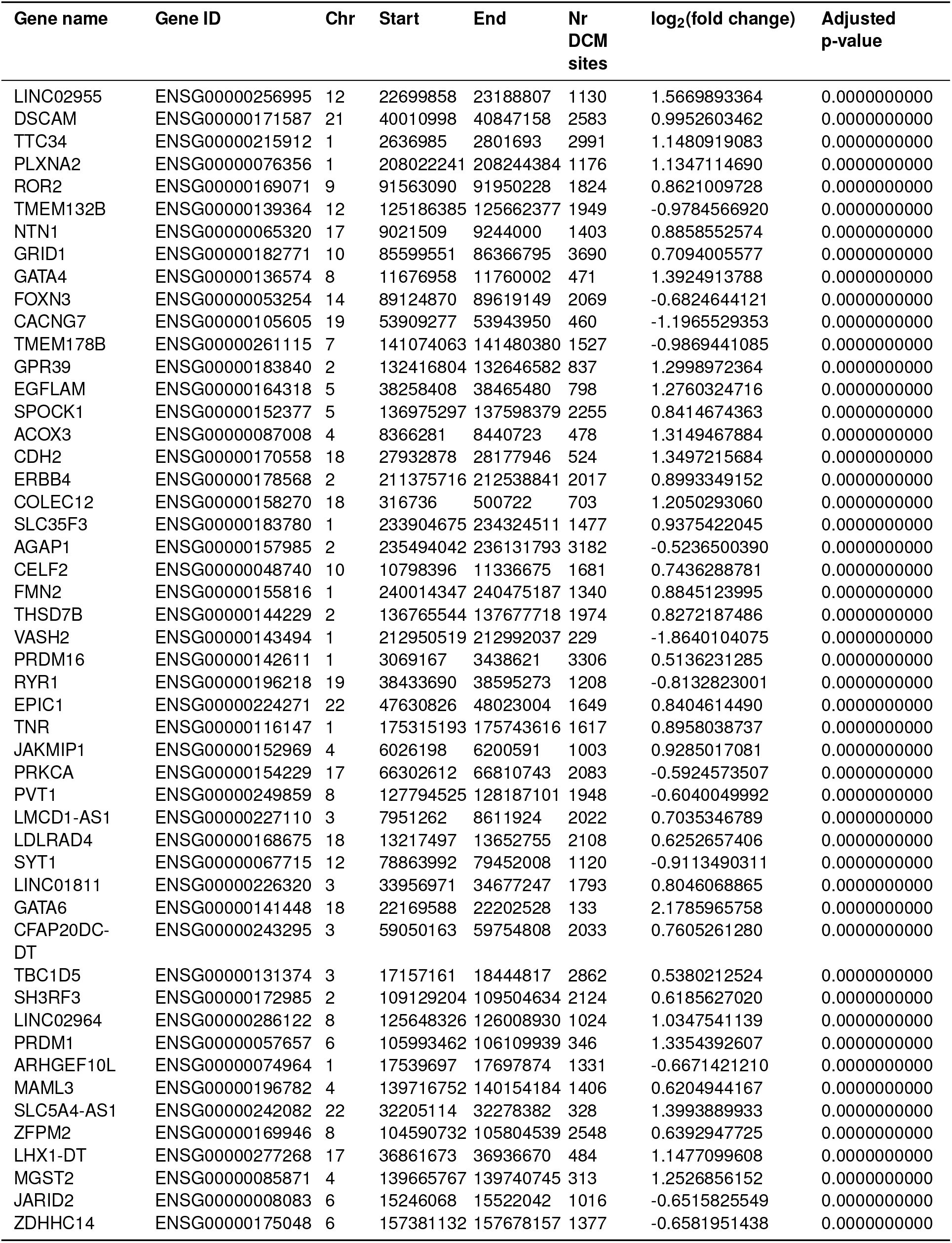
Differentially DCM-methylated genes between iPSC^dox^ and DE^dox^. Only the most significant genes are shown.

**Table S5.**
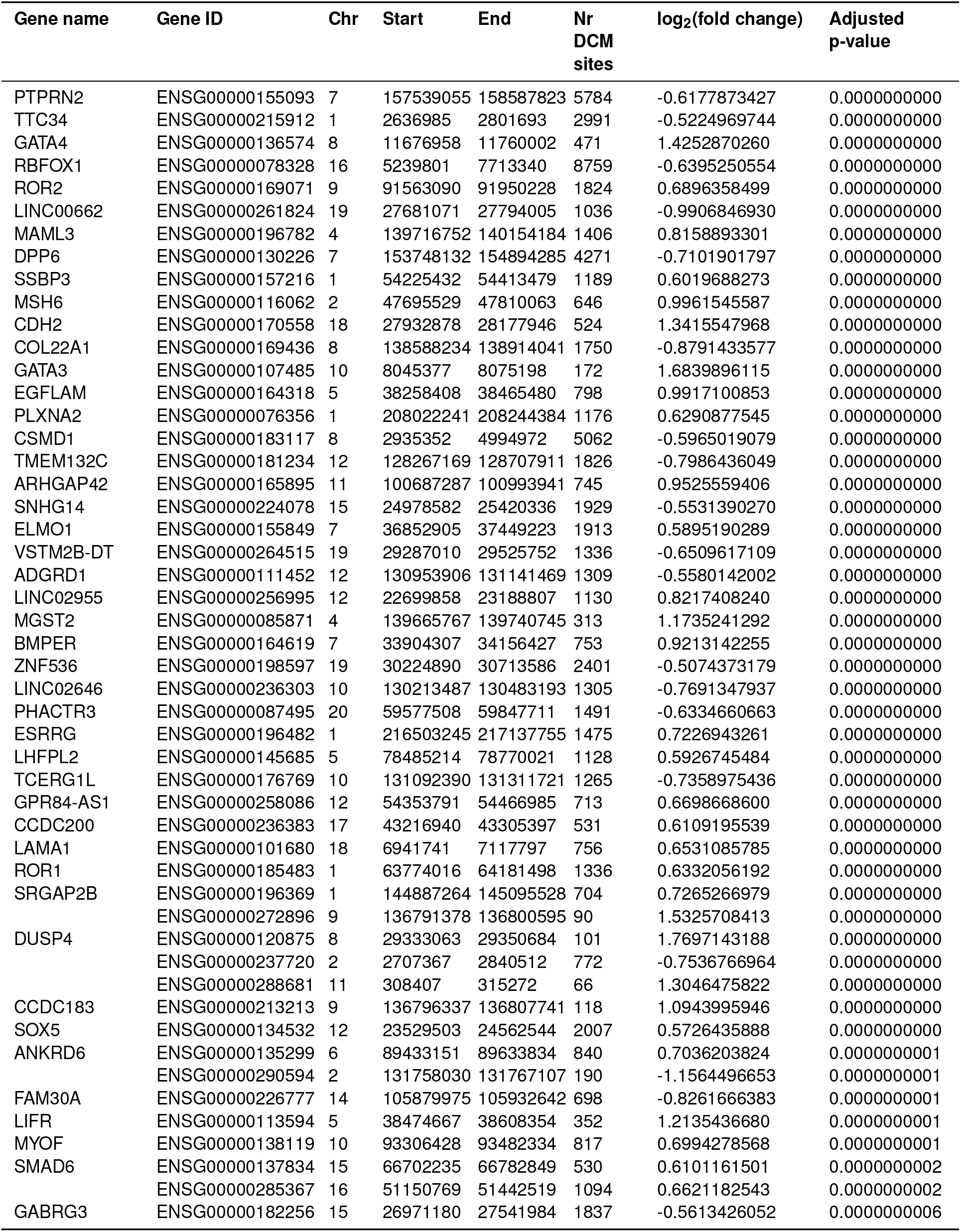
Differentially DCM-methylated genes between DE^chase^ and PGT^chase^. Only the most significant genes are shown.

